# Modulating the Blood-Brain Barrier by Light Stimulation of Molecular-Targeted Nanoparticles

**DOI:** 10.1101/2020.10.05.326843

**Authors:** Xiaoqing Li, Vamsidhara Vemireddy, Qi Cai, Hejian Xiong, Peiyuan Kang, Xiuying Li, Monica Giannotta, Heather Hayenga, Edward Pan, Shashank Sirsi, Celine Mateo, David Kleinfeld, Chris Greene, Matthew Campbell, Elisabetta Dejana, Robert Bachoo, Zhenpeng Qin

**Author notes:** These authors contributed equally to this work (Xiaoqing Li, Vamsidhara Vemireddy, Qi Cai).

## Abstract

The blood-brain barrier (BBB) tightly regulates the entry of molecules into the brain by tight junctions that seals the paracellular space and receptor-mediated transcytosis. It remains elusive to selectively modulate these mechanisms and to overcome BBB without significant neurotoxicity. Here we report that light stimulation of tight junction-targeted plasmonic nanoparticles selectively opens up the paracellular route to allow diffusion through the compromised tight junction and into the brain parenchyma. The BBB modulation does not impair vascular dynamics and associated neurovascular coupling, or cause significant neural injury. It further allows antibody and adeno-associated virus delivery into local brain regions. This novel method offers the first evidence of selectively modulating BBB tight junctions and opens new avenues for therapeutic interventions in the central nervous system.

**One Sentence Summary:** Gentle stimulation of molecular-targeted nanoparticles selectively opens up the paracellular pathway and allows macromolecules and gene therapy vectors into the brain.

---

The burden of neurodegenerative diseases continues to rise and the development of effective therapies has slowed in part due to the challenges of the blood-brain barrier (BBB) (*1-4*). As a highly selective barrier, the BBB restricts penetration of molecules and particles from blood to the brain parenchyma by two main features, including the tight junction (TJ) complex that seals the paracellular space between the endothelial cells, and low level of transcytosis through the endothelium (*5-8*). Prior approaches to circumvent the BBB have led to limited success: convection enhanced delivery has limited success in the clinical trials (*9*), intrathecal delivery has limited parenchymal penetration (*10*). Methods to increase BBB permeability such as using high osmotic mannitol have significant neurotoxicity (*11, 12*). Focused ultrasound stimulation of circulating gas microbubbles stretches the blood vessels to permeate the BBB, but can lead to vascular spasm (*13*), neuroinflammation (*14*) and disruptions in brain connectivity (*15*). It remains unknown whether a gentle perturbation could temporarily modulate the BBB permeability without substantial vasculature or brain injury.

Laser excitation of gold nanoparticles (AuNPs) has been extensively studied for photothermal cancer therapy, relying on the heat generation of AuNPs to raise the tumor temperature to above physiological temperatures (*16, 17*). However, short laser pulse excitation of AuNPs leads to different phenomena, including photoacoustic effect that involves heating water next to AuNPs under a nanosecond laser, and mechanical wave generation due to AuNP expansion from picosecond (ps) or other short laser pulses (*16, 18, 19*). We developed a new and simple method to modulate the BBB with ps laser excitation of TJ-targeted plasmonic AuNPs (Fig. 1A).

**Fig. 1.**
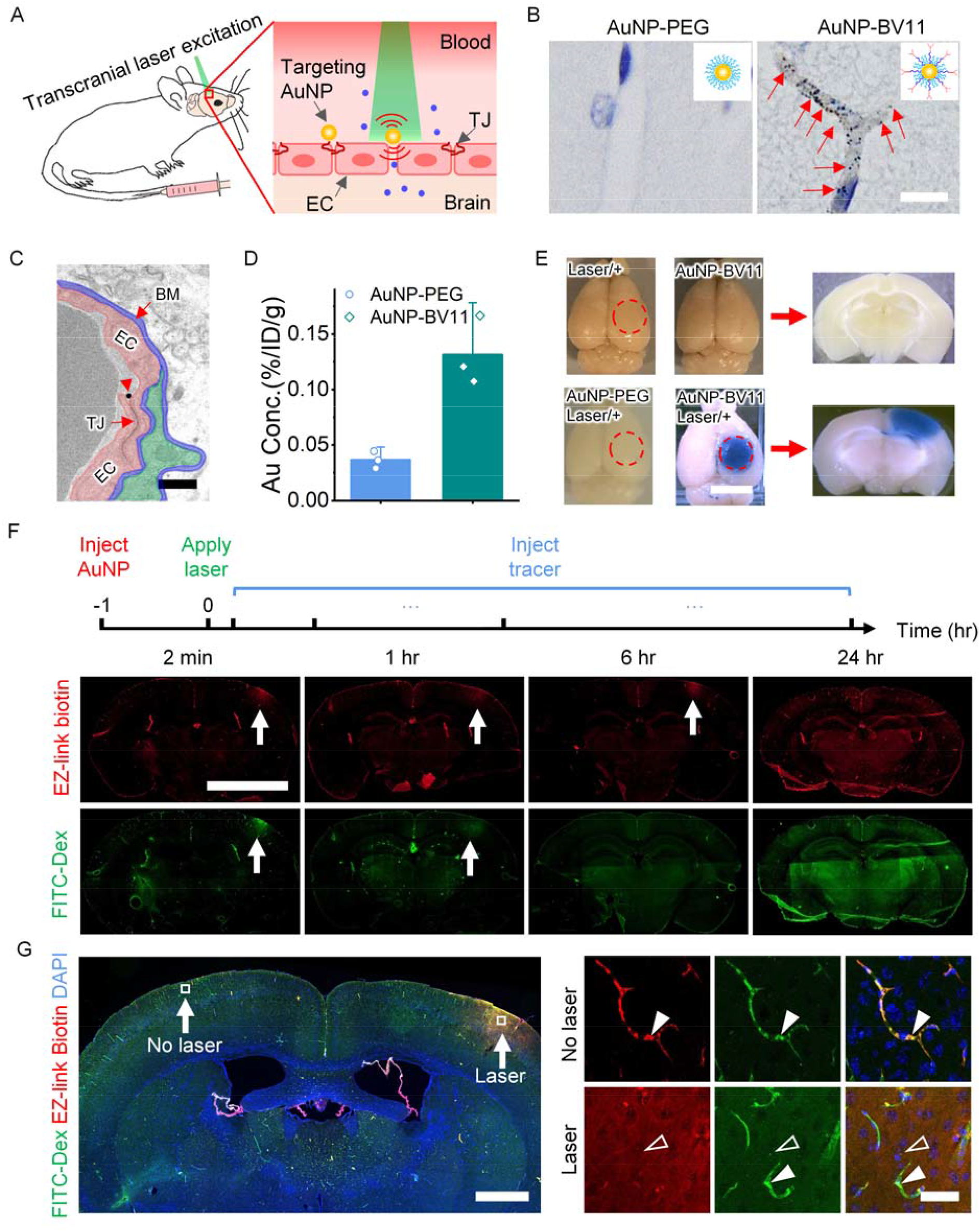
Light stimulation of TJ-targeting nanoparticles reversibly modulates BBB permeability. (A) Schematic for transcranial laser stimulation of TJ-targeting AuNP (AuNP-BV11) for BBB modulation. AuNP-BV11 interacts with ps laser to perturb the TJ to modulate the BBB permeability. Blue dots represent molecules penetrating into the brain. (B) AuNPs are visualized by silver enhancement staining in the brain. (C) AuNP-BV11 (arrow head) co-localizes with TJ detected by electron microscopy (EM). Pseudocolors: endothelial cell (EC: red), basement membrane (BM: blue), pericyte (P: green), endothelial cell (EC, pink). (D) Quantification of AuNP-BV11 and AuNP-PEG accumulation in the brain. (E) BBB modulation visualized by the leakage of albumin-binding Evans blue (25 mJ/cm^2^,1 pulse). (F) BBB permeability probed by molecular tracers (660 Da EZ-link biotin and 70kDa FITC-dextran (FITC-Dex), 5 mJ/cm^2^, 1 pulse). (G) Left panel: brain coronal section shows dyes leakage at laser excitation region in contrast to contralateral side with no laser treatment. Right panels: inserts in left panel show both EZ-link biotin and FITC-dextran were confined to the blood vessels (arrow head) without laser treatment, while the laser excitation of AuNP-BV11 caused EZ-link biotin and FITC-dextran leak into the brain parenchyma (empty arrow head) (5 mJ/cm^2^, 1 pulse, 2 minutes post laser treatment). Scale bar: 10 µm (B), 400 nm (C), 4 mm [(E), (F)], 1 mm (G), 40 µm (G insert). Data expressed as Mean ± SD (n=3).

## Nanoparticle targeting and light modulation of the BBB

First, we tested the feasibility to target the tight junction. AuNPs were synthesized and conjugated with the antibody BV11 (AuNP-BV11) (Fig. S1), to target a TJ component, junctional adhesion molecule A (JAM-A). JAM-A is expressed on the luminal (blood-facing) side of the BBB (*20*). We examined the biodistribution and targeting of AuNP-BV11 with intravenous (IV) injections in mice (Fig. 1A). Silver enhancement staining shows AuNP accumulation along cerebral vessels only in the case of AuNP-BV11, but not for non-targeting AuNP-PEG (PEG: polyethylene glycol) (Fig. 1B). Electron microscopy (EM) imaging shows that AuNP-BV11 co-localizes with the TJ (Fig. 1C), indicating that AuNP-BV11 recognizes and targets TJ *in vivo*. AuNP-BV11 shows 4 times higher concentration in the brain than AuNP-PEG (Fig. 1D), much shorter circulation time (t1/2=10 minutes for AuNP-BV11 versus t1/2=2.3 hours for AuNP-PEG), and dramatically different organ distributions (Fig. S2A-E). Moreover, IV administration of AuNP-BV11 doesn’t cause obvious toxicity as observed by measuring body weight and hematoxylin and eosin (H&E) staining of major organs (Fig. S2F and S3).

Laser excitation of the TJ-targeted AuNP-BV11 temporarily increases BBB permeability, as indicated by the Evans blue (EB) leakage into the brain tissue including grey matter and white matter, while laser alone, or AuNP-BV11 injection alone, or laser excitation of AuNP-PEG does not lead to the BBB modulation (Fig. 1E). We administered tracers with different molecular weight to detect the time window of leakage. The 660 Da EZ-link biotin was detected up to 6 hours, while 70 kDa fluorescein isothiocyanate (FITC) labeled dextran and albumin-binding EB (66 kDa total) were only detected up to 1 hour after laser excitation (5 mJ/cm^2^, 1 pulse, Fig. 1F and S4-7 and S8A-C). This size-dependent leakage at 6 hours suggests a paracellular pathway with a compromised TJ between endothelial cells. The tracer leakage is further demonstrated by confocal imaging (Fig. 1G). No leakage of the 3 tracers was observed 24 hours after laser excitation, suggesting that the BBB is recovered within this time period. Furthermore, the depth of BBB modulation (1 to 2.9 mm) is dependent on laser fluence, consistent with Monte Carlo simulation of light propagation in mouse brain (Fig. S8D. The laser beam size can be conveniently tuned to control the area of BBB modulation (Fig. S8E) and a laser fiber modulates the BBB in deeper brain regions (Fig. S8F).

## Impact of BBB modulation on vascular dynamics and brain parenchyma

Arteriole vasomotion is the dilation and constriction of arterioles to meet energy demands in the brain and forms the basis of blood-oxygen-level-dependent imaging (BOLD) in functional magnetic resonance imaging (fMRI) (*21*). Changes in the vasomotion can impair brain’s ability to respond to energy demands and thus neurovascular coupling. To examine whether BBB modulation impairs vasomotion, we imaged blood vessels using two-photon *in vivo* imaging in awake, head-fixed mice before and after laser stimulation (Fig. 2A-C, 5 mJ/cm^2^, 1 pulse). Fourier transform analysis suggests that the arteriole vasomotion (centered around 0.1 Hz) was persistent before and after BBB modulation (from 1 to 24 hours, Fig. 2D). As expected, venules do not show vasomotion (Fig. 2E). These results suggest that BBB modulation doesn’t impair vasomotion and associated neurovascular coupling.

**Fig. 2.**
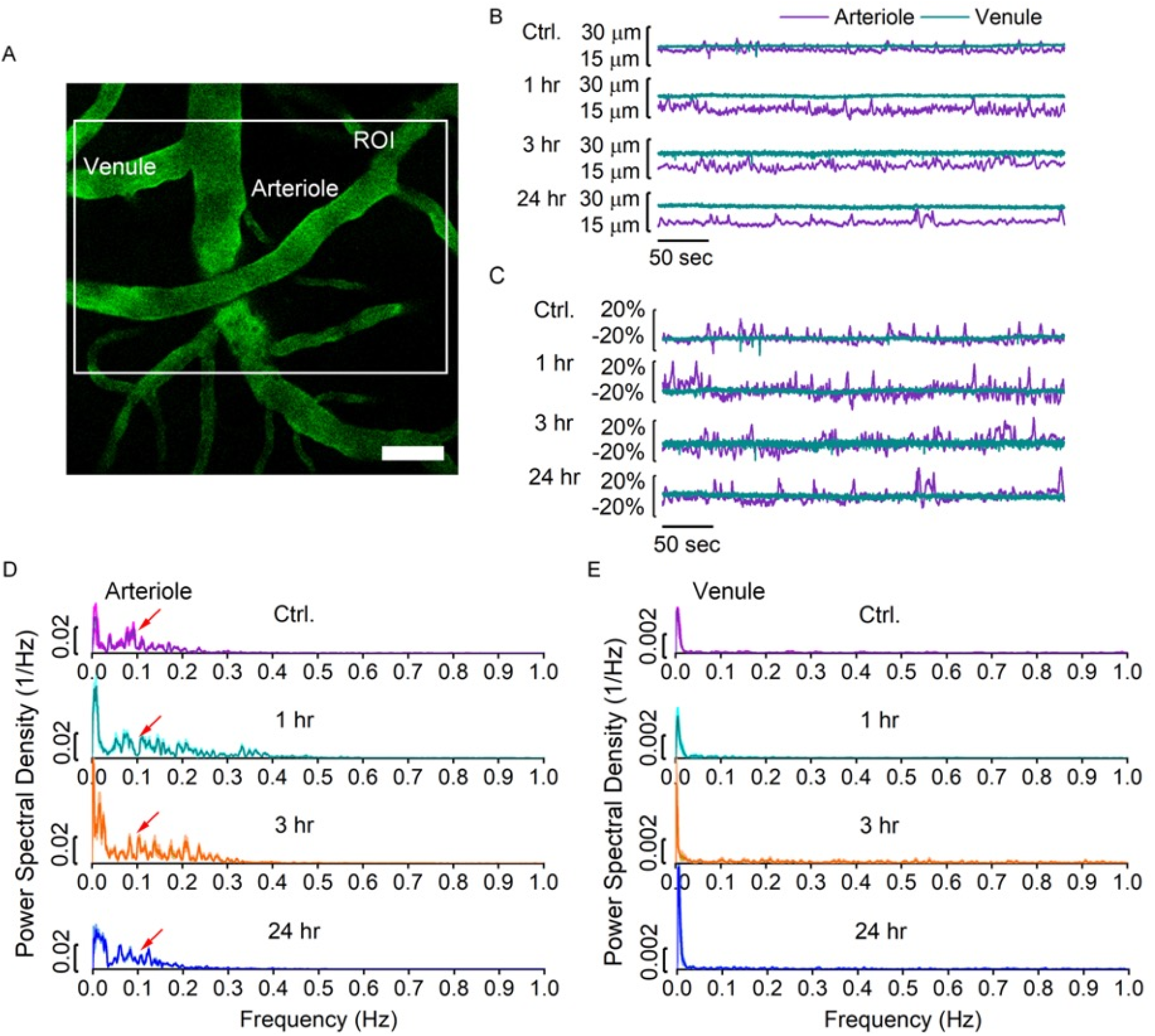
BBB modulation does not disrupt spontaneous vasomotion in the awake mouse. (A) Representative *in vivo* two photon microscopy image of a fluorescent angiogram through a thinned-skull window in an awake, head-fixed mouse. White box indicates a selected ROI containing a pair of an arteriole and a venule. (B-C) The diameter changes and the percentage change of diameter of the arteriole and venule segment over the recorded time course (5 mJ/cm^2^, 1 pulse). (D-E) Fourier transform analysis of the percentage change of diameter in (C), demonstrating the arteriole oscillations around 0.1 Hz (D, indicate by the arrows). No spontaneous vasomotion in venule segment was observed (E). Shaded areas represent SEM. Scale bar, 50 µm.

We further investigated the impact of BBB modulation on the brain vasculature and parenchyma. No significant changes in cerebral vascular density was observed by labelling the vessels with lectin and glucose transporter Glut1 (Fig. S9). Observation of the brain ultrastructure showed no obvious abnormalities in endothelial cell (EC), basement membrane (BM) and pericyte morphology (Fig. S10). Enlarged astrocyte endfeet were observed at 0.5 and 6 hours post laser stimulation (Fig. S10), suggesting the BBB modulation may cause water influx into the brain and lead to uptake by astrocytes endfeet (*22*). The synapses and mitochondria appeared intact (Fig. S10). Moreover, the Golgi staining shows that BBB modulation preserves dendritic spines of neurons (Fig. S11). Further analysis shows no significant changes in neuronal nuclues and axon (NeuN, Ankyrin-G), water transporter of astrocyte endfeet (AQP4), and pericyte (NG2) (Fig. S12). We observed an increase of Iba1^+^ microglia and GFAP^+^ astrocyte with heterogeneity (Fig. S13-14). Previous studies showed the critical role of GFAP^+^ astrocyte in BBB repair in injury or diseased conditions (*23-26*), which suggests that astrocytes may engage in the BBB recovery after laser stimulation.

## BBB modulation by paracellular diffusion

Next, we examined the route of BBB modulation by laser excitation of plasmonic AuNPs. Low levels of fluid phase transcytosis and the presence of TJ are two main factors that limit the permeability of the BBB. We performed transcardial perfusion with 2% lanthanum nitrate after BBB modulation for electron microscopy (EM) imaging (Fig. S15A-B). The EM results reveal TJs in 51% capillaries are completely filled with lanthanum (La) and TJs in 49% capillaries show incomplete infusion with lanthanum with laser treatment (Fig. 3A). In contrast, all TJs show incomplete infusion with lanthanum in control mouse without laser treatment (Fig. 3A). Importantly for the laser treated brain, lanthanum diffused into the basement membrane and interstitial space (Fig. 3B and S15C). On the other hand, no lanthanum tracers were observed in the intracellular vesicles. This result provides a clear evidence that BBB modulation involves the paracellular pathway to allow tracer to pass through the TJ into basement membrane and brain interstitial space.

**Fig. 3.**
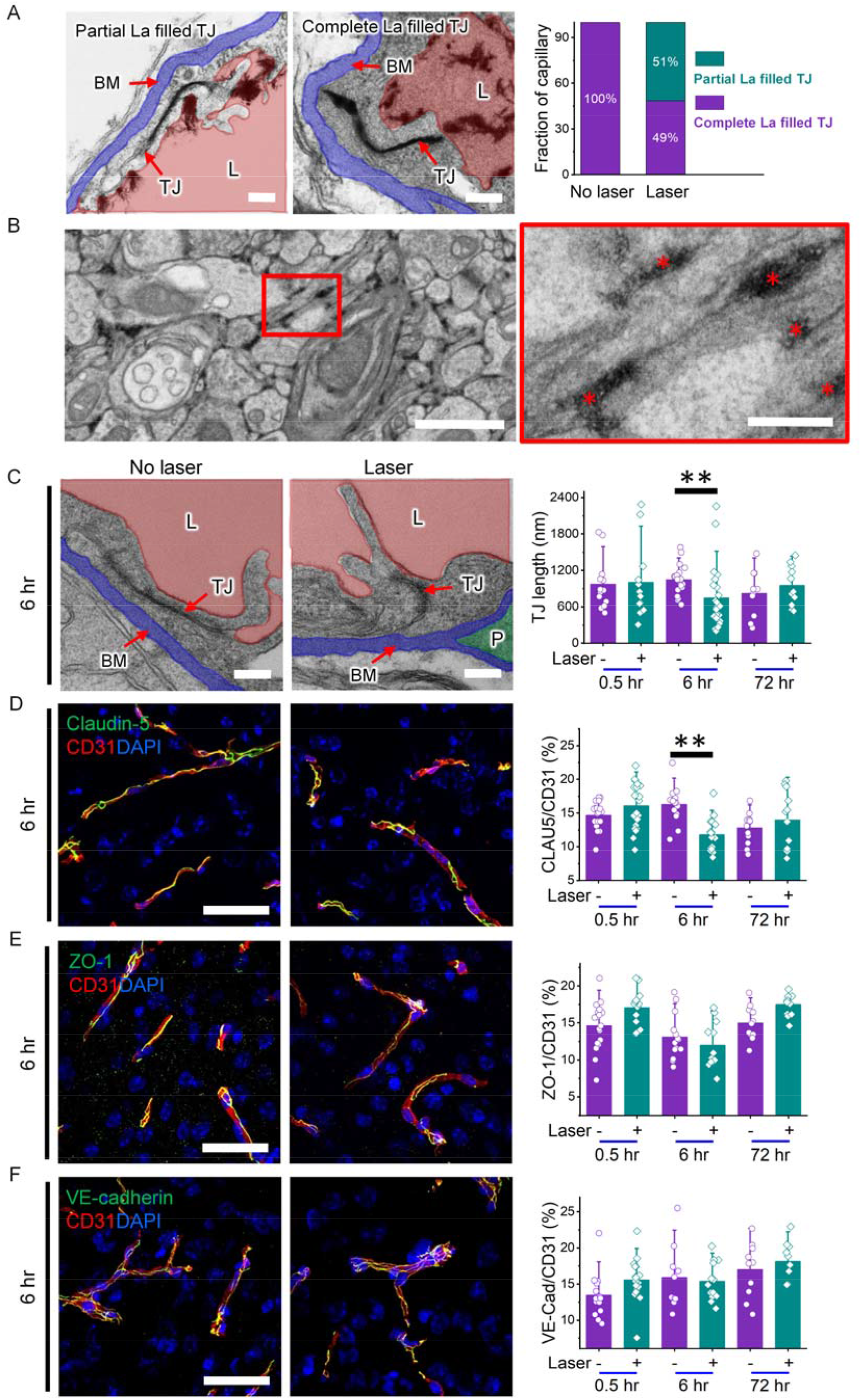
BBB modulation involves paracellular route and compromised tight junctions. (A) Electron microscopy imaging of lanthanum (La) infused brain microvasculature (25 mJ/cm^2^,1 pulse, 2 hours post laser treatment). Glycocalyx is visible on the lumen wall. (B) Electron microscopy imaging of La diffusion into brain interstitial space under same condition, labeled by asterisk (*) in the zoomed in picture. (C) Electron microscopy imaging (5 mJ/cm^2^,1 pulse, 6 hours post laser) and quantification of TJ length at three time points. (D-F) Immunohistochemical staining of TJ protein claudin-5 (D), TJ-associated protein ZO-1 (E) and adherens junction protein VE-cadherin (F). Tight junction (TJ), lumen (L: red), basement membrane (BM: blue), pericyte (P). Scale bars: 200 nm [(A), (B insert), (C)], 1 µm (B), 40 µm [(D), (E), (F)]. Data expressed as Mean ± SD (n>15). ** p < 0.05.

Furthermore, EM imaging reveals that the TJ length decreased before returning to the same as control at 72 hours (Fig. 3C). We then stained junctional proteins and found that claudin-5 in the TJ complex decreased at 6 hours and recovered at 72 hours (Fig. 3D), while the TJ-associated protein ZO-1 and the adherens junction protein VE-cadherin didn’t change significantly (Fig. 3E-F). These results confirm the involvement of the paracellular pathway and the change of claudin-5 in BBB modulation.

## Gene and antibody delivery to the brain

We further examined the ability to delivery gene therapy vectors and antibody by BBB modulation. Gene therapy using adeno-associated viruses (AAV) and antibodies has shown promise for a number of neurological diseases such as Parkinson disease and aromatic-L-amino acid decarboxylase (AADC) deficiency (*27*). However, one central challenge is the delivery of AAV into targeted cells in the body. We injected AAV systemically along with AuNP-BV11 and applied laser stimulation. The result shows that AAV crossed the BBB and infected 64% of neurons at right hemisphere with laser illumination, compared to the left hemisphere with no virus crossing the barrier (Fig. 4A-C). Although some AAV serotypes have shown capability to cross the BBB and shows widespread transfection in the nervous system (*28*), our approach allows delivery of gene therapy vectors in local brain regions. Furthermore, we detected endogenous mouse IgG and systemically-injected human IgG after BBB modulation (Fig. 4D). The gene and antibody penetration into the brain indicates potential therapeutic opportunities with BBB modulation.

**Fig. 4.**
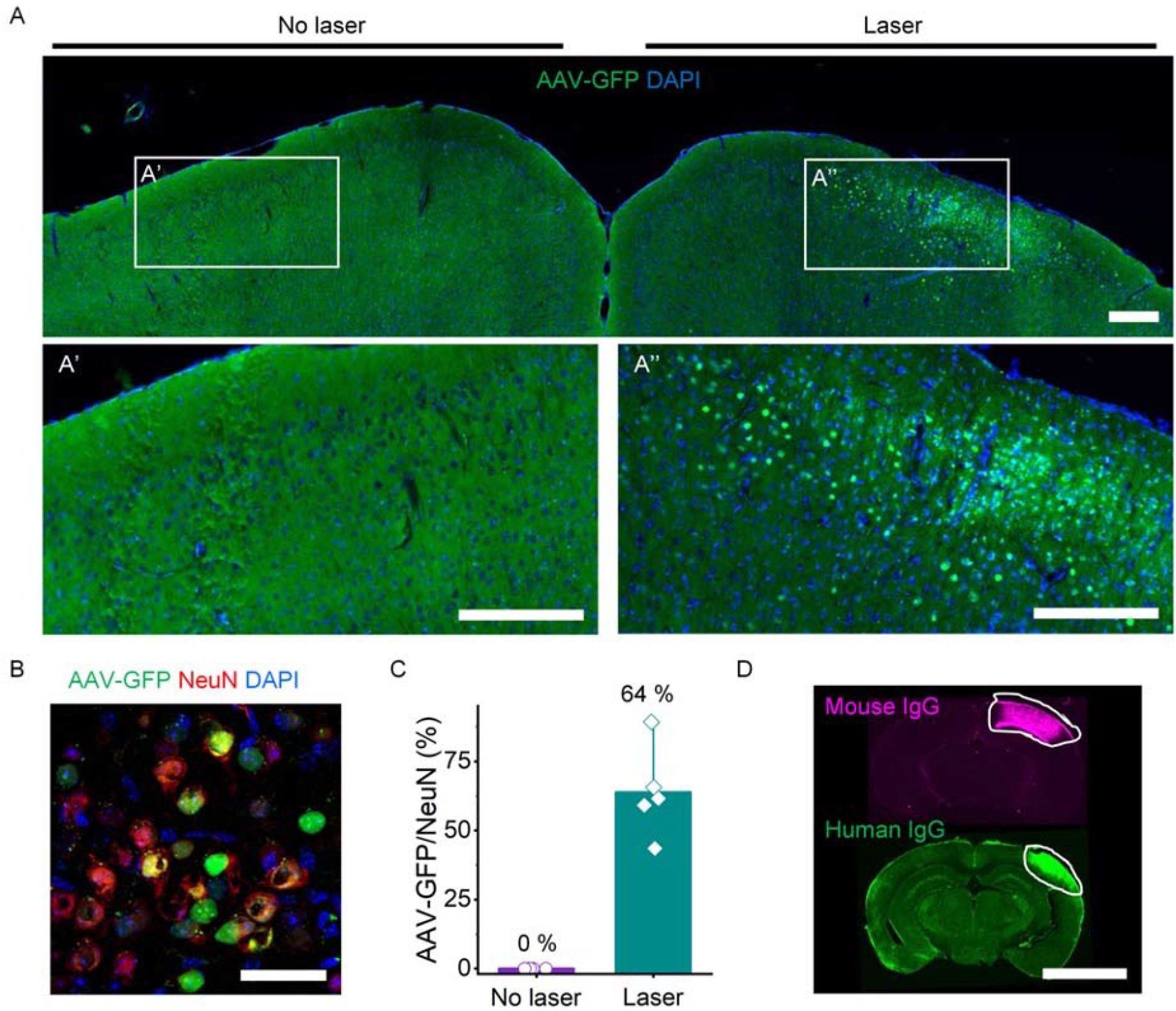
BBB modulation enables delivery of gene therapy vector and IgG antibody. (A) Delivery of AAV-CamKII-GFP into the brain (10 mJ/cm^2^, 1 pulse). Right: laser excitation of TJ-targeting AuNPs; left: no laser. (B) Colocalization of AAV-GFP and NeuN. (C) Percent of neuron (NeuN) expressing GFP. (D) Antibody delivery into the brain by BBB modulation. Scale bar: 200 µm [(A), (A’), (A’’)], 40 µm (B), 4 mm (D). Data expressed as Mean ± SD (n=5).

## Impact of BBB modulation on neural stem cell niche

Lastly, we investigated the impact of BBB modulation on perivascular microenvironment that are critical for many important processes in the brain including neural stem cell niche. The ventricular-subventricular zone (V-SVZ) (Fig. S16A) is one of the neural stem cell niches and provides neuronal precursors to migrate to olfactory bulb and differentiate into interneurons each day in adult mice (*29, 30*). The vascular system in V-SVZ regulates the neurogenesis by the spatial cues and signals (*31*). Changes in the V-SVZ microenvironment can impact the proliferation of neural stem cells. Using transgenic nestin::GFP mice, we observed the nestin^+^ neural stem cells interact intimately with blood vessels in the V-SVZ region (Fig. S16B). Ki67 indicates the actively dividing neural stem cells (Fig. S17B). BBB modulation was evidenced by the dye leakage in the V-SVZ zone on the right hemisphere (Fig. S16C). The proliferating cells (ki67^+^ and BrdU^+^) in V-SVZ region decreased significantly at 72 hours post laser stimulation. The changes in neural stem cell proliferation suggests that it is sensitive to changes in the BBB microenvironment (Fig. S16D-F). Active modulation of BBB changes the perivascular microenvironment, and can prove useful to investigate a wide range of processes that rely on brain vasculature such as gliomagenesis (*32*) and cancer cell migration (*33*).

## Summary

The BBB restricts molecular penetration into the brain by suppressing transcytosis and promoting tight junction formation and integrity. So far, no studies have demonstrated the ability to selectively and safely modulate specific mechanisms in local brain regions. Light stimulation of TJ-targeted AuNPs introduces specific changes in TJ to allow macromolecules and virus particles to cross the BBB. Increase in BBB permeability is temporary and recovers within hours. Close examination of brain microvasculature and parenchyma indicates no disruptions in vasodynamics or neuronal injury. Active modulation of BBB alters the perivascular microenvironment for neural stem cells and may lead to new understanding of cancer and stem cell biology in the brain (*34*). Further research combining synthetic biology may open up new opportunities for neuroscience research (*35*). Focal delivery of genes and antibodies will provide insight into the treatment of neurological disorders (*36, 37*).

## Acknowledgments

The authors thank the Histo Pathology Core at University of Texas Southwestern Medical Center (UTSW) for assistance with histology staining, Dr. Bret Evers for assistance with H&E analysis, the Electron Microscopy Core of UTSW for assistance with EM samples process, Yaning Liu and Haihang Ye for assistance with EM imaging of gold nanoparticles, Stephanie Shiers for assistance with NeuN and Ankyrin-G staining, Susanne J van Veluw and Steven S Hou for providing code of vasomotion analysis and members of the Qin laboratory for discussions.

## Author contributions

Xiaoqing Li, Vamsidhara Vemireddy, and Qi Cai designed and performed the experiments and contributed equally to this work. Hejian Xiong participated in biodistribution study, tail vein injection and brain tissue processing. Peiyuan Kang performed the simulation. Xiuying Li participated in initiating the research idea. Monica Giannotta and Elisabetta developed the anti-JAM-A antibodies for this study. Heather Hayenga, Shashank Sirsi and Edward Pan participated in discussion and provided suggestions. Celine Mateo and David Kleinfeld developed the protocol for vasomotion study. Chris Greene and Matthew Campbell developed the protocol for tight junction staining and EZ-link biotin staining. Robert Bachoo and Zhenpeng Qin supervised the project. All authors provided critical feedback and helped shape the research, analysis and manuscript.

## Data and materials availability

All data is available in the main text or the supplementary materials.

