## Supplemental Materials for "Modulating the Blood-Brain Barrier by Light Stimulation of Molecular-Targeted Nanoparticles"

**This PDF file includes:**

Materials and Methods
Figs. S1 to S16
Table S1
References (38-48)

### Materials and Methods

#### Materials

Anti-JAM-A antibody BV11 was provided by Drs Elisabeth Dejana and Monica Giannotta at FIRC Institute of Molecular Oncology Foundation. Gold (III) chloride, fluorescein isothiocyanate-labeled dextran (FITC-dextran), Evans blue, 4′,6-diamidino-2-phenylindole (DAPI), EZ-link biotin, DMSO, hydroquinone, sodium citrate tribasic, tween 20, triton-X 100, sucrose, horseradish peroxidase (HRP), human Immunoglobulin G (IgG), Evans blue and Lanthanum (III) nitrate hexahydrate were purchase from Sigma-Aldrich. OPSS-PEG-SVA, mPEG-thiol were purchased from Laysan Bio, Inc. Li silver enhancement kit, donkey serum, goat serum, DyLight 594 labeled tomato lectin, gold reference standard solution, phosphate buffered saline, borate buffer, syringes, needles, 20 kDa MWCO dialysis membrane were purchased from Thermo Fisher Scientific. All chemicals were analytical grade. Adult mice C57BL/6 mice (7 weeks old, 22-25 g) were ordered from Charles River Laboratories. Animal protocols were approved by Institutional Animal Care Use Committee (IACUC) of University of Texas at Dallas.

#### Gold nanoparticle conjugation with antibodies

Gold nanoparticles (AuNPs) were synthesized following previously reported method (*38*) and conjuated with antibodies (Fig. S1A). Briefly, the size of AuNPs used in this work is 50 nm confirmed by transmission electron microscopy (TEM) (Fig. S1B). BV11, an anti-JAM-A antibody, was diluted to 0.5 mg/ml in PBS, followed by dilution in 2 mM borate buffer at pH8.5 to 0.05mg/ml. OPSS-PEG-SVA was dissolved in borate buffer and quickly added to the diluted antibody at a 125:1 molar ratio. The mixture was vortexed briefly and kept for 3 hours shaking on ice, followed by dialysis at 4 °C overnight to remove free OPSS-PEG-SVA through 20 kDa MWCO membrane. The thiolated antibodies were reacted with concentrated AuNPs at a 200:1 molar ratio for 1 hour on ice. To stabilize AuNP-BV11, mPEG-thiol was added at 6 PEG/nm$^2$ for backfilling (*39*) the empty space of AuNPs for 1 hour on ice. Finally, the modified AuNPs were washed 3 times and characterized by dynamic light scattering (DLS) and UV-vis spectroscopy.

#### AuNP biodistribution *in vivo*

To quantitatively measure AuNP distribution *in vivo*, we perfused the animal and collected major organs at 1, 6, 12 and 24 hours after I.V. injection of AuNPs. The tissue was then digested in aqua regia and centrifuged at 5 000 g for 5 minutes. The gold concentration was analyzed by inductively coupled plasma mass spectrometry (ICP-MS).

To microscopically examine AuNP-BV11 distribution, major organs were excised for histology staining using silver enhancement regents. We performed paraffin processing of tissues with 20 µm thick, followed by deparaffined step. The sections, then, were rinsed by Milli-Q water and incubated with Li silver enhancement developer for 20 minutes at room temperature. After washing in water, the treated tissues were counterstained with hematoxylin for 5 minutes, followed by dehydration through 100% ethanol and clearance by xylenes. Finally, the tissues were covered with synthetic mounting media and imaged under bright field microscopy.

*In vivo* toxicity measurement of TJ-targeting AuNPs

To investigate the dose safety of TJ-targeting AuNPs (AuNP-BV11), we weighed the animals to check any body weight loss after I.V. injections of 36.9 µg/g nanoparticle and 100 µl saline as control group. We further performed hematoxylin and eosin (H&E) staining to observe any pathological changes in major organs by comparing the nanoparticle injected group with control animal group. We injected AuNP-BV11 (36.9 µg/g) and saline (100 µl) intravenously. After 23 days, mice were slowly transcardially perfused with PBS and 4% paraformaldehyde (PFA). Brain and other organs were extracted and post-fixed with 4% PFA at -4 °C for 12 hours. The Histo Pathology Core at University of Texas Southwestern Medical Center assisted in preparing H&E staining. Imaging was performed with Slide Scanner (VS120, Olympus).

Optical modulation of the BBB *in vivo* and data analysis

C57BL/6 mice were used for the *in vivo* experiments. The animal was anesthetized by 2-3% isoflurane (in air) and intravenously administrated 36.9 µg/g AuNPs functionalized by anti-JAM-A antibody BV11. 1% lidocaine mixed with 0.5% bupivacaine was subcutaneously injected on the scalp. Then, the scalp was peeled back to expose the skull. Finally, the picosecond (ps) laser was applied through intact skull to excite the AuNPs in blood vessels. Tracers, EZ-link biotin (2 mg/ml in saline, 100 µl) and FITC-dextran (40 mg/ml in saline, 100 µl), or Evans blue (2% in PBS, 100 µl) were injected into tail vein at different time points post laser stimulation. After 0.5 hours, mice were transcardially perfused with 30 ml PBS to remove the intravascular dye, followed by 4% PFA. The brain and other organs were extracted for imaging and histology staining.

To quantify the intensity of tracers, fixed brains were sliced 40 µm thick at -20 °C on cryostat. We picked up brain coronal sections by glass slides and imaged by Slide Scanner (VS120, Olympus). Samples, injected with tracers EZ-link biotin and FITC-dextran, were incubated with Cy3-labelled streptavidin (1:200) to detect EZ-link biotin and DAPI to label nuclei, followed by imaging. Fluorescent intensity was analyzed by FIJI imageJ for each brain section. For each mouse, total fluorescent intensity was summed up from individual sections.

To study the delivery of gene therapy vector, AAV-9 (AAV.CamKII.HI.GFP-Cre.WPRE.SV40) was injected into the tail vein of a mouse treated with laser excitation of AuNP-BV11. After 7 days, the animal was perfused with icy PBS, followed by 4% PFA. Post-fixing was performed with 4% PFA at 4 °C for 12 hours. The brain tissue was then dehydrated in 30% sucrose and sliced 20 µm thick at -20 °C on cryostat. Free-floating coronal sections were stained with NeuN for investigating the percentage of neurons expressing AAV-GFP.

Light propagation simulation

We firstly calculated the light fluence profile with an infinitesimal point light source by Monte Carlo Multilayered (MCML) program developed by Wang et al (*40*). Then, to obtain the finite beam light propagation profiles, the point source results were used for convolution over a Gaussian beam profile with program in Matlab 2016a. The optical properties used in the MCML program were obtained from previous studies (*41-44*) and were listed in Table S1.

We built the geometry of brain in the simulation according to the mouse brain (*45*). There are five layers in the model. The first layer is skull with thickness of 0.3 mm. The meninges were laid in between of skull and brain with a thickness of 0.1 mm. Because mouse meninges are very thin and contains mostly liquid, we use cerebrospinal fluid (CSF) properties for this layer in the simulation. The cortex is grey matter with thickness of 1 mm, followed a layer of white matter (corpus callosum) with thickness of 0.3 mm. The thickness of bottom grey layer is greater than 4 mm. The laser beam profile can be described by Gaussian pulse formula:

$$G(r) = \frac{2P}{\pi R^2} \exp\left(-2\frac{r^2}{R^2}\right)$$

where R is the $1/e^2$ diameter of 4.1 mm (Full width at half maximum = 2.9 mm), which was obtained from blade edge measurements. Briefly, a blade was used to cover the beam gradually and pulse energy was measured simultaneously. By fitting the data with Gaussian beam model, the beam size was obtained (*46*). P is the total energy of single laser pulse irradiation (0.03 mJ). The results were analyzed by Matlab 2016a.

Immunohistochemistry staining

To immunostain tight junction proteins, Glut1, AQP4 and NG2, the mouse brain tissue was snap-frozen on dry ice once quickly removed from the skull and stored at -20 °C before processed 20 μm thick on cryostat. The coronal brain sections were fixed for 10 minutes using icy methanol at -20 °C. Blocking solution (5% normal donkey serum, 0.1% triton X-100 in PBS) was applied to the tissue for 1 hour at room temperature. After washing, the sections were incubated with primary antibodies first (diluted ratio 1:500): rabbit anti-claudin-5 (341600, Fisher Scientific), rabbit anti-ZO-1(402200, Fisher Scientific), rabbit anti-VE-cadherin (361900, Fisher Scientific), rabbit anti-Glut1 (RB9052, Fisher Scientific), mouse anti-AQP4 (SC-32739, Santa Cruz) and rabbit anti-NG2 (AB5320, Millipore), overnight at 4°C and then secondary antibodies 2 hours at room temperature, followed by incubation with DAPI solution.

To stain Iba1, GFAP, ki67 and doublecortin (DCX), the mouse brains were dehydrated in 30% sucrose solution. The dehydrated brains were sliced 20 μm thick on a freezing cryostat. Coronal sections were washed in PBS for 3 times to remove cryoprotectant solution completely. After blocking (20% normal goat or donkey serum with 0.1% Triton X-100 in PBS) for 1 hour at room temperature, free-floating sections were incubated with primary antibodies: goat anti-Iba1 (AB48004, Abcam), rabbit anti-GFAP (RB087A0, Fisher Scientific), rabbit anti-ki67 (RM9106, Fisher Scientific) and rabbit anti-DCX (4604, Cell Signaling) for two nights at 4°C. Secondary antibodies then were added, followed by incubation with DAPI solution.

For staining of NeuN and Ankyrin-G, the mice were perfused with PBS and 4% PFA at 5 ml/min, followed by post-fixing in 4% PFA for 1 hour. We sliced the brain tissue with thickness of 30 μm on cryostat at -20 °C after dehydration in 30% sucrose solution. The coronal brain sections were washed in milli-Q water at 37 °C for 5 minutes, and incubated with pepsin solution for antigen retrieval. The treated sections were then incubated with blocking buffer at room temperature for 1 hour, and primary antibodies: rabbit anti-NeuN (NC1284461, Fisher Scientific) and mouse anti-Ankyrin-G (75146, Antibodies Incorporated), overnight at 4°C. Secondary antibodies were applied for 1 hour at room temperature, followed by incubation with DAPI solution.

To study the effect of optical BBB modulation on the neural stem cell niche, we applied a laser with flux of 48 mJ/cm$^2$ on the right hemisphere for deep BBB modulation at ventricular-subventricular zone (V-SVZ). The BrdU (B5002, Sigma) were injected at 4 hours and 2 hours, respectively, before perfusion with PBS and 4% PFA. The brain tissue was dehydrated in 30% sucrose, and post fixed in 4% PFA. We stained ki67 as previous description (the same protocol as the staining of Iba1) with primary antibodies and secondary antibodies incubation. After washing in PBS, the sections were then incubated with 2M HCl with 0.1% triton-X-100 for 10 minutes at room temperature and treated in 0.1 M borate buffer, followed by incubation with mouse anti-BrdU antibodies (347580, BD Biosciences) and secondary antibodies. Nuclei were stained by DAPI solution.

All the treated sections were mounted to glass slides carefully and imaged by a Slide Scanner (VS120, Olympus), or confocal microscopy (FV3000RS, Olympus), or Spinning Disk microscopy (SD-OSR, Olympus).

The following secondary antibodies were used and purchased from Fisher Scientific: donkey anti-mouse IgG alexa 488 (A21202); donkey anti-mouse IgG alexa 594 (A21203); donkey anti-mouse IgG alexa 647 (A31571); donkey anti-rabbit IgG Alexa 488 (A21206,); donkey anti-rabbit IgG Alexa 594 (A21207); donkey anti-rabbit IgG Alexa 647 (A31573); donkey anti-goat IgG AFP 488 (A11055); goat anti-mouse IgG AFP 488 (PIA32723); goat anti-rat IgG Alexa 594 (A11007); goat anti-mouse IgG1 555 (A21127); goat anti-chicken IgG AFP 488 (A11039).

Transmission electron microscopy (TEM)

To image the morphology of AuNPs, we put AuNPs solution on a copper grid and the samples were dry at room temperature.

To study the brain ultrastructure post laser stimulation, the mice were transcardially perfused with icy PBS, followed by a fixative solution containing 2% PFA and 2% glutaraldehyde. Post-fixing was carried out in same fixing solution for 1 day. We then sliced the brain tissue including the regions with and without laser stimulation. The Electron Microscopy Core at University of Texas Southwestern Medical Center assisted in preparing the TEM samples for imaging.

To study the pathways of BBB modulation, the mice were transcardially perfused with 2% lanthanum (La) in PBS following icy PBS perfusion, then perfused by 2% PFA and 2% glutaraldehyde. Post-fixing was carried out in same fixing solution for 1 day. We then sliced the brain tissue including the regions with laser stimulation indicated by Evans blue leakage and the regions without laser treatment. The Electron Microscopy Core at University of Texas Southwestern Medical Center assisted in preparing the TEM samples for imaging.

TEM imaging was carried out by TEM-JEOL 1400+ (JEOL).

Image analysis

All images were analyzed by FIJI imageJ. To study the changes of junctional proteins, area fraction of claudin-5, ZO-1 and VE-cadherin was obtained and normalized by CD31 indicating cerebral vessel. Vasculature density was analyzed by area fraction of lectin. Area fraction of Glut1, AQP4 was normalized by CD31 indicating blood vessels. Area fraction of NG2 and GFAP was normalized by DAPI. Counts of NeuN, Ankyrin-G and Iba1 per field were normalized by counts of DAPI.

Thinned-skull window surgical method

Thinned-skull window surgery was performed as described previously (*47*). Mice were anesthetized with 5% isoflurane (in air) and placed in a stereotactic frame equipped with a gas anesthesia mask (Kopf instruments). Isoflurane was maintained at 1.5%-2% throughout the surgical procedure. The rectal temperature was monitored using a feedback temperature probe and a heat pad (DC Temperature controller, FHC). The skull was exposed and a thin layer of cyanoacrylate glue was applied to the entire skull surface. A square area of the skull (2×2 mm, 1 mm lateral and 1 mm posterior to bregma) was carefully thinned to 20 µm with a hand-held drill (EXL-M40, Osada). A thin layer of cyanoacrylate glue was applied over the window, and a cover glass (12-542-C, Fisher Scientific, cut to 2 mm in length) was secured to the window. A custom-made stainless-steel head bar was attached to the skull using cyanoacrylate glue and dental cement (C&B Metabond® Quick Adhesive Cement System, Parkell). Mice received one subcutaneous injection of buprenorphine (0.05 mg/kg) post-surgery and were allowed to recover for 3-4 days before imaging.

Two-photon microscopy set-up

Mice were imaged using a FVMPE-RS two-photon laser-scanning system (Olympus). An INSIGHT DS-OL laser system generated two-photon excitation at 840 nm to record vascular activity. External detectors containing a photomultiplier tube (PMT) collected emitted light in the range of 495-540 nm (green). PMT settings were kept constant for each experiment, but laser power was adjusted slightly as needed. 40x dipping water immersion objective (Olympus) was applied in the vasomotion recordings.

Vasomotion recordings

Half an hour before imaging, 0.1 ml of 2000 KDa FITC-dextran dye (25 mg/ml) dissolved in sterile PBS was administered with retro-orbital injection under brief anesthesia to label the blood flow. The mice were stabilized on a head restraint apparatus and were fully awake. Spontaneous vasomotion of was recorded in arteriolar and venules (identified based on their morphology and direction of blood flow) at 100 µm below the pial surface. The video was recorded continuously over 6-7 minutes at a speed of 2 µs/pixel, a resolution of 1.6091 pixel/micron, and a frame-rate of 10 Hz (4000 frames were recorded). The control video was collected first before ps laser stimulation, and the same animal was used to collect videos after ps laser stimulation (5 mJ/cm$^2$, 1 pulse) at 1, 3 and 24 hours post laser irradiation. 2-3 ROIs were imaged per mouse. The mice were returned to their home cage between imaging sessions for food and water.

Data analysis for vasomotion

Recorded videos were analyzed with FIJI imageJ and MATLAB (version R2019a) using custom-written scripts. Videos were first aligned to correct for motion in the x-y plane using a plugin. Vessel diameter (D) was determined using Full Width at Half maximum (FWHM) method as reported in literature (*21*). Percentage change of vessel diameter was calculated by the following equation:

$$\%\,change = \frac{D_{frame} - D_{average}}{D_{average}} \times 100\%.$$

A discrete Fourier transform was performed for each percentage vessel diameter trace and the vasomotion was calculated in the range between 0-1 Hz (*48*).

Statistical Analysis

All the data were plotted and analyzed with Origin 2020. For the *in vivo* biodistribution study using ICP-MS, one-way ANOVA was used to compare the accumulation difference in main organs between targeting and non-targeting gold nanoparticles. For the kinetics study using different tracers, one-way ANOVA was used to compare the total fluorescent intensity between different time points. To study the depth of the BBB modulation, one-way ANOVA was used to compare the depth among various laser fluence. We performed one-way ANOVA to analyze confocal and TEM images of brain tissue to compare the difference between laser and no laser treatment.

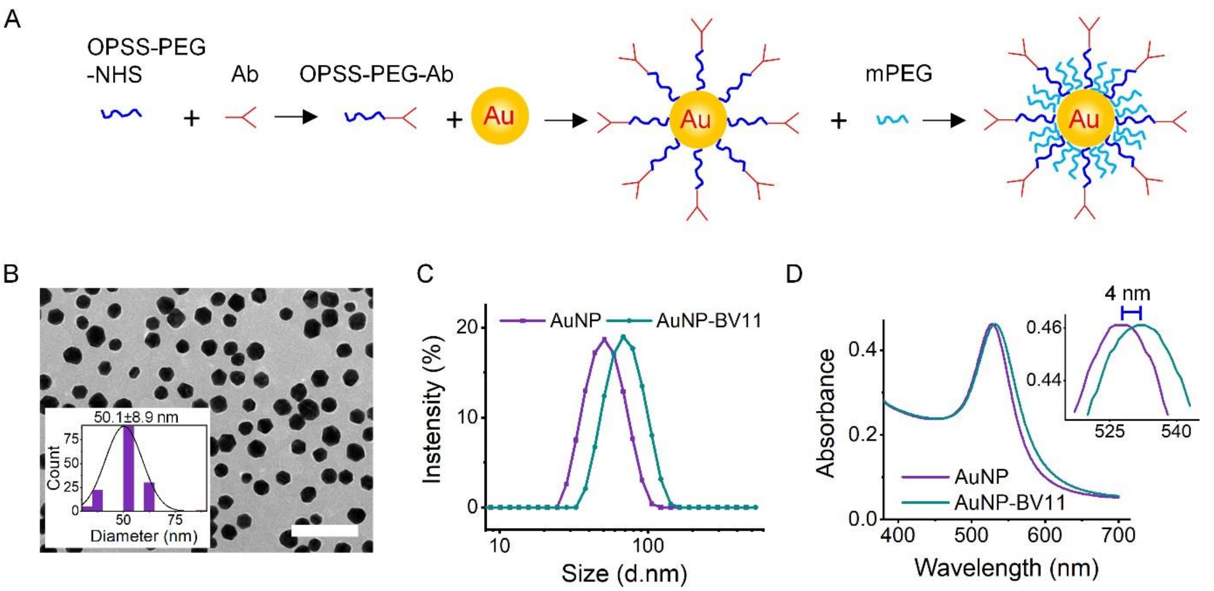


**Fig. S1. Synthesis and characterization of TJ-targeting AuNPs (AuNP-BV11)**. (A) The process of AuNP conjugated with anti-JAM-A antibody (Ab, BV11). (B) TEM image of AuNPs. Insert is the size distribution histogram of AuNPs (n=95). (C) The size increases by 20 nm after conjugation measured by DLS. (D) The absorbance peak shifts by 4 nm after conjugation measured by UV-vis spectroscopy. Scale bar: 200 nm.

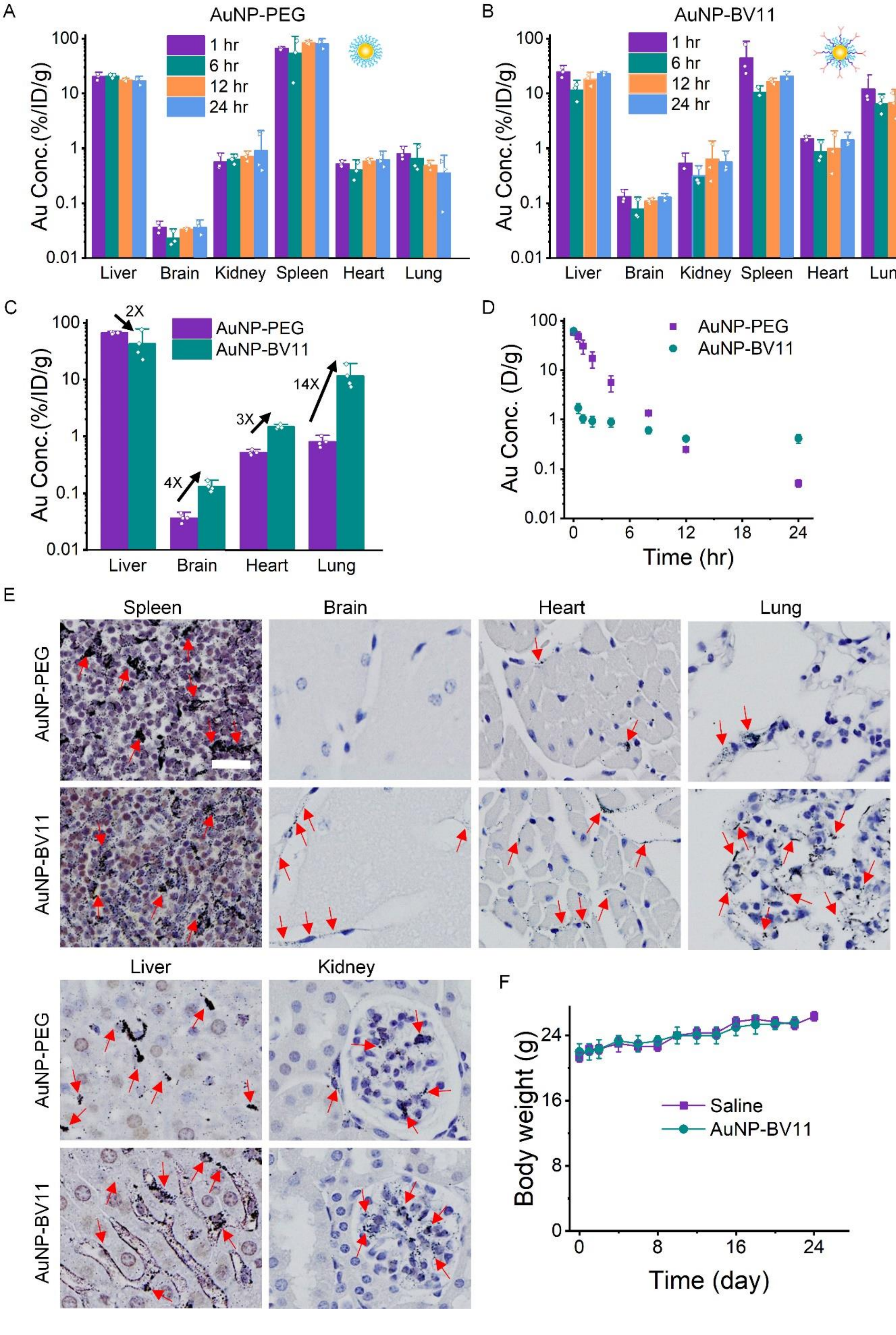
A
AuNP-PEG
1 hr
6 hr
12 hr
24 hr
Au Conc.(%/ID/g)
Liver
Brain
Kidney
Spleen
Heart
Lung
B
AuNP-BV11
C
2X
4X
3X
14X
AuNP-PEG
AuNP-BV11
D
Au Conc. (D/g)
Time (hr)
E
Spleen
Brain
Heart
Lung
Liver
Kidney
F
Saline
AuNP-BV11
Body weight (g)
Time (day)

**Fig. S2.** ***In vivo*** **biodistribution and toxicity study of TJ-targeting AuNPs**. (A-B) ICP-MS analysis of AuNP-PEG (A) and AuNP-BV11 (B) distribution in major organs, %ID/g: % injection dose/gram. (C) Comparison of biodistribution between AuNP-PEG and AuNP-BV11 at 1 hour post injection. (D) Concentration of AuNP-PEG and AuNP-BV11 in blood stream. $T_{1/2}$ is10 minutes for AuNP-BV11, and it is 2.3 hours for AuNP-PEG. (E) Silver enhancement of AuNPs in major organs. (F) Body weight for saline and AuNP-BV11 injected animals. Doses of AuNP-PEG and AuNP-BV11 are 36.9 µg/g. Saline: 100 µl. Scale bar: 20 µm. Data expressed as Mean ± SD (n =3).

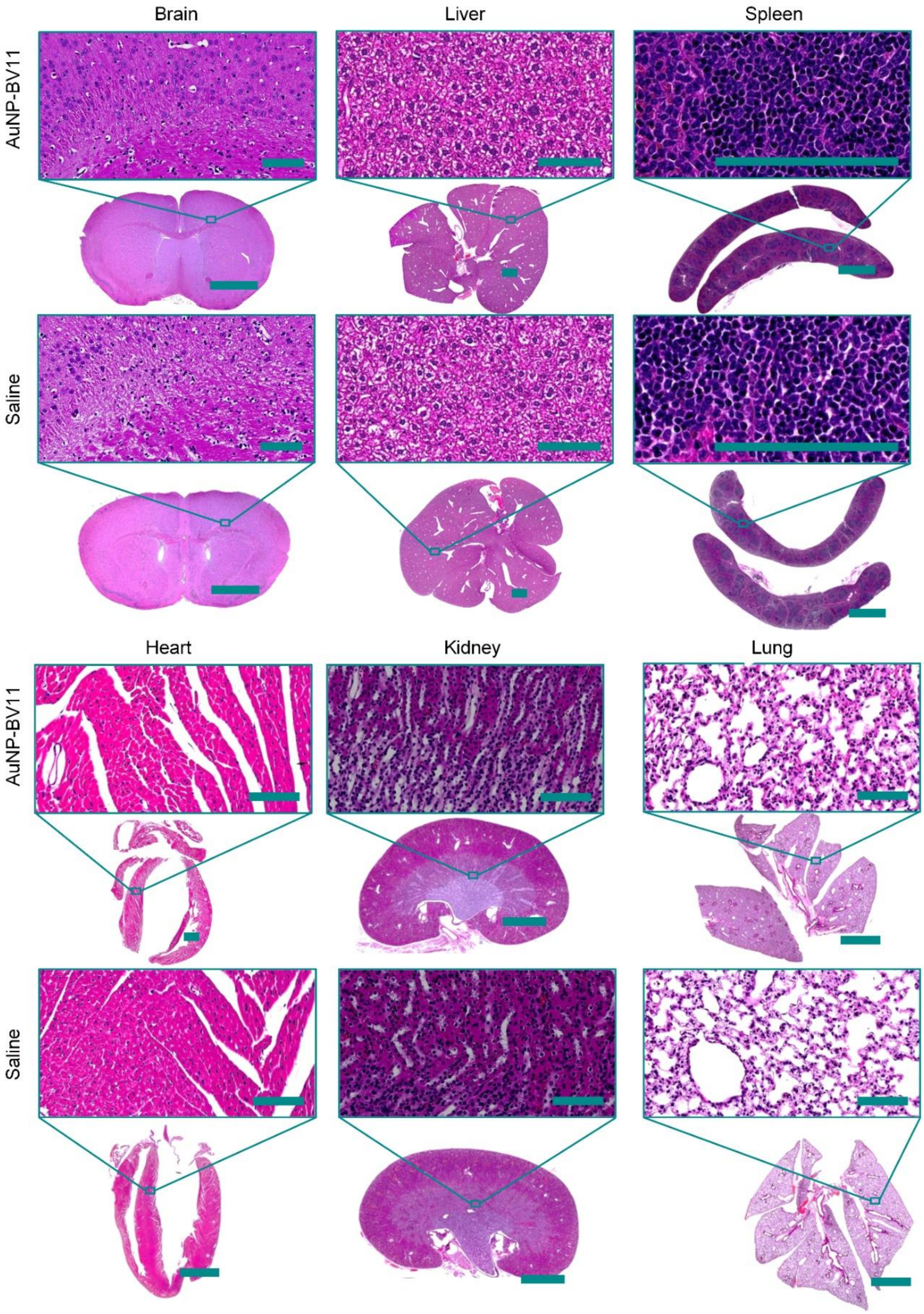


**Fig. S3. H&E staining in main organs for AuNP-BV11 and saline injected animals**. AuNP-BV11: 36.9 µg/g. Saline as control: 100 µl. Scale bar: 2 mm in low magnification, 100 µm in high magnification.

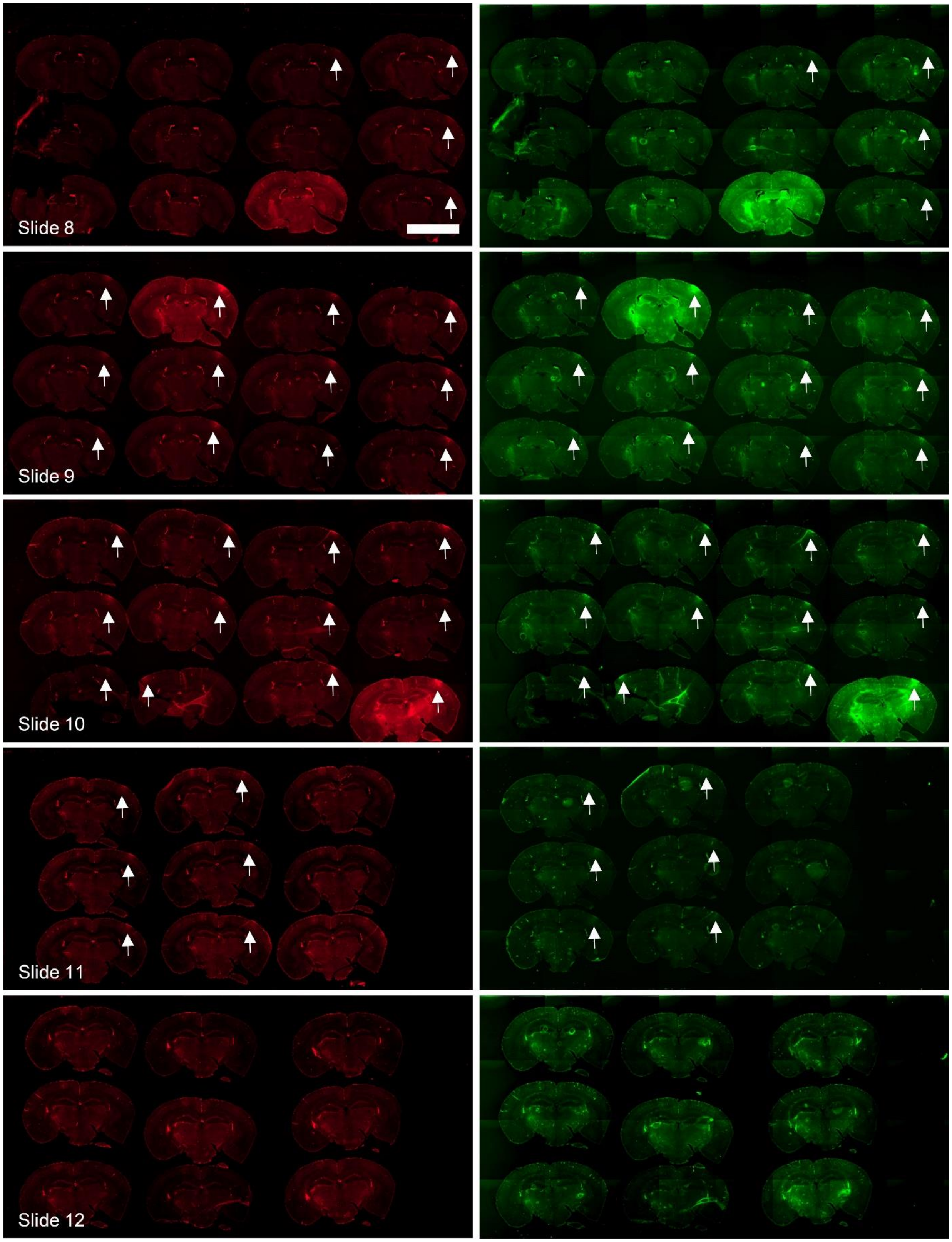


**Fig. S4. Small and large molecule extravasation 2 minutes post laser excitation (5 mJ/cm$^2$, 1 pulse).** Red: 660 Da EZ-link biotin. Green: 70 kDa FITC-dextran. Arrow: signal of tracers. Scale bar: 5 mm.

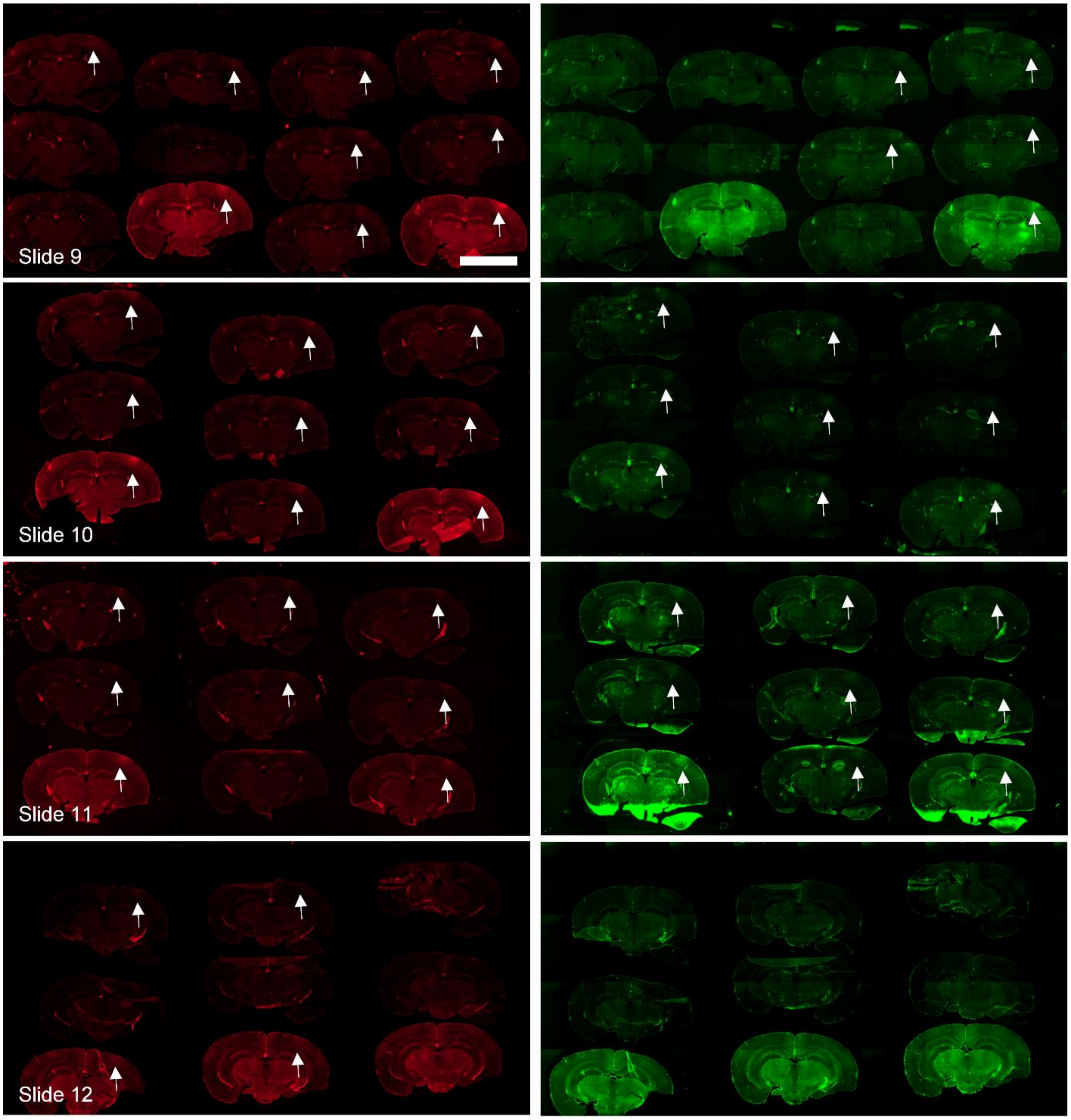


**Fig. S5. Small and large molecule extravasation 1 hour post laser excitation (5 mJ/cm$^2$, 1 pulse).** Red: 660 Da EZ-link biotin. Green: 70 kDa FITC-dextran. Arrow: signal of tracers. Scale bar: 5 mm.

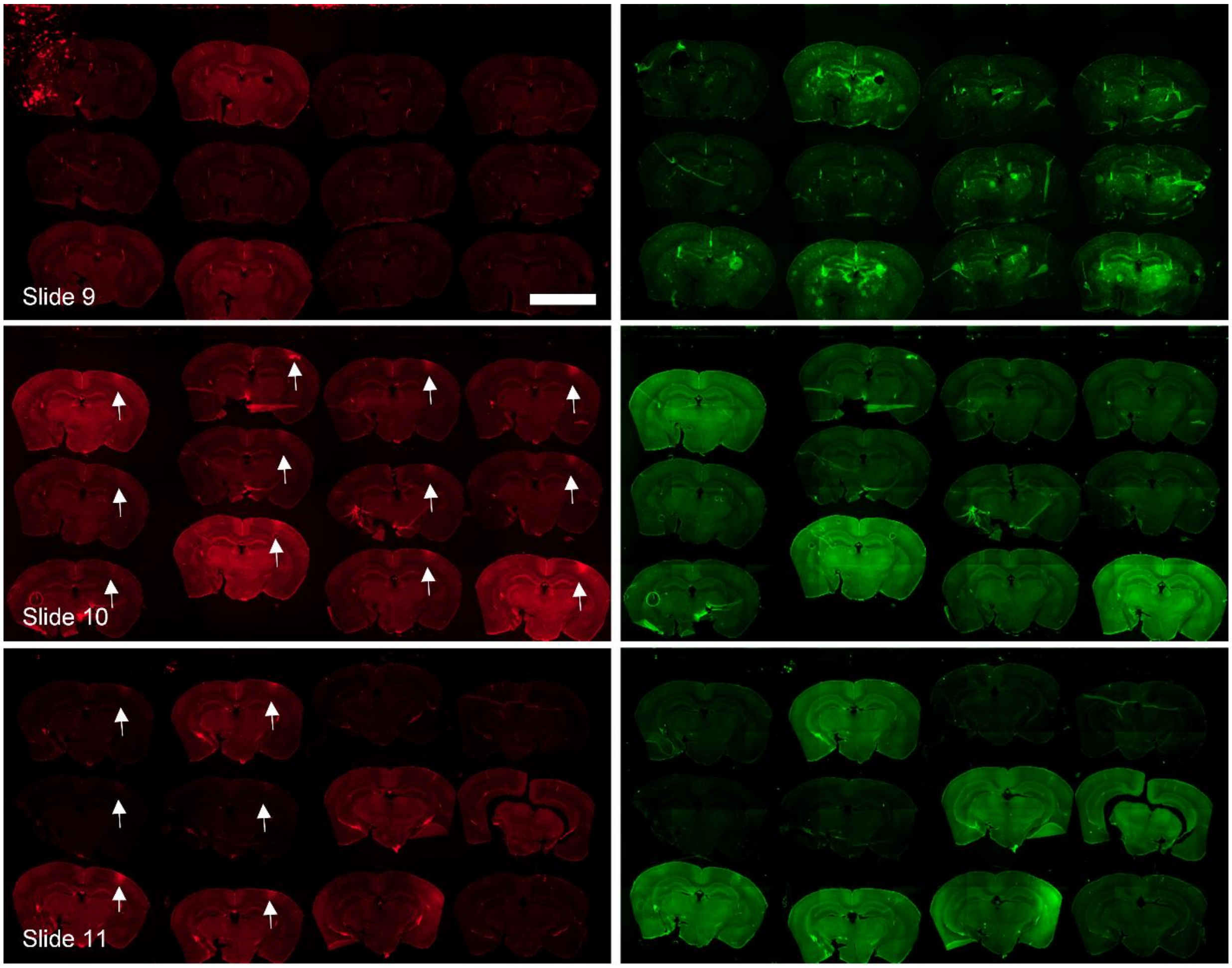


**Fig. S6. Small molecule extravasation 6 hours post laser excitation (5 mJ/cm$^2$, 1 pulse).** Red: 660 Da EZ-link biotin. Green: 70 kDa FITC-dextran. Arrow: signal of tracers. Scale bar: 5 mm.

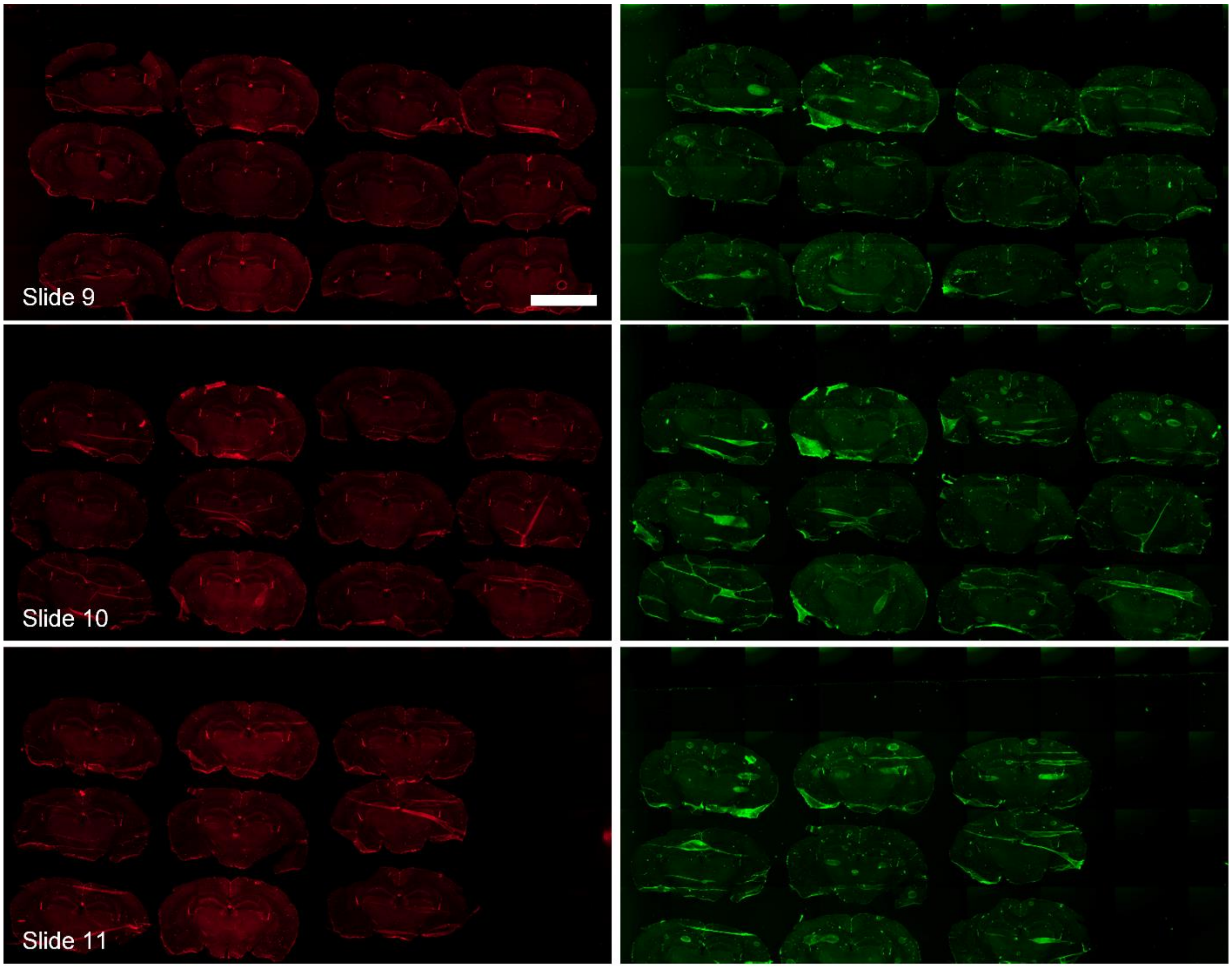


**Fig. S7. No extravasation 24 hours post laser excitation (5 mJ/cm$^2$, 1 pulse).** Red: 660 Da EZ-link biotin. Green: 70 kDa FITC-dextran. Scale bar: 5 mm.

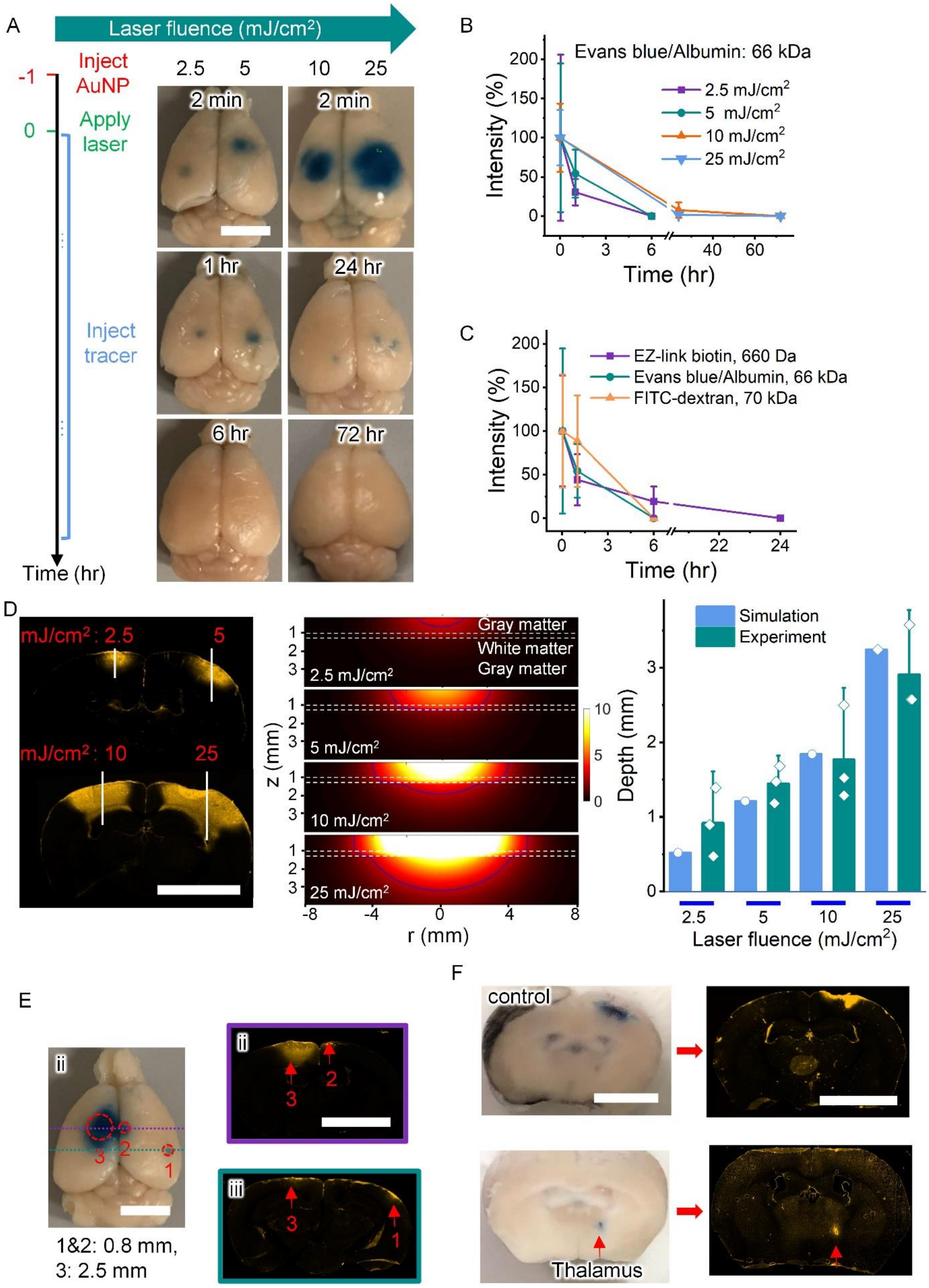
A
Laser fluence (mJ/cm²)
Inject AuNP
Apply laser
Inject tracer
Time (hr)
2.5 5 10 25
2 min
1 hr
24 hr
6 hr
72 hr
B
Evans blue/Albumin: 66 kDa
2.5 mJ/cm²
5 mJ/cm²
10 mJ/cm²
25 mJ/cm²
Intensity (%)
Time (hr)
C
EZ-link biotin, 660 Da
Evans blue/Albumin, 66 kDa
FITC-dextran, 70 kDa
D
mJ/cm² : 2.5
mJ/cm² : 10
Gray matter
White matter
2.5 mJ/cm²
5 mJ/cm²
10 mJ/cm²
25 mJ/cm²
z (mm)
r (mm)
Simulation
Experiment
Depth (mm)
Laser fluence (mJ/cm²)
E
1&2: 0.8 mm,
3: 2.5 mm
F
control
Thalamus

**Fig. S8. Reversible and precise BBB modulation *in vivo*.** (A) Albumin-binding Evans blue (EB) leakage into the brain under different laser fluence (2.5-25 mJ/cm$^2$, 1 pulse) and at different times after laser exposure. (B) Quantification of EB extravasation with various laser fluence. (C) Kinetics of tracer leakage by quantifying the samples with different tracers (from Fig. S4-7 and S8A) (5 mJ/cm$^2$, 1 pulse). Small molecule 660 Da EZ-link biotin was detected up to 6 hours, while large molecules (66 kDa EB/albumin, 70 kDa FITC-dextran) were detected only up to 1 hour. (D) Depth of BBB modulation (left: Evans blue fluorescence, middle: Monte Carlo simulation of laser fluence distribution in mouse brain; right: comparison of experiment and simulation). Blue lines in the middle panel represent the threshold fluence to modulate BBB predicted by Monte Carlo simulation. (E) Localized BBB modulation by manipulating laser beam size (25 mJ/cm$^2$, 1 pulse). The BBB modulation is indicated by EB signal which was detected by fluorescent microscopy. Diameter of beam size: 0.8 mm for spots 1 and 2, 2.5 mm for spot 3. (F) Precise BBB modulation in thalamus via optical fiber with core diameter 200 µm to deliver light (10 mJ/cm$^2$, 5 pulses, 5 Hz). Upper panels: only insert optical fiber to reach the thalamus without laser delivery. Bottom panels: optical fiber carrier light for modulating the BBB in thalamus. Scale bar: 4 mm. Data expressed as Mean $\pm$ SD (n =3).

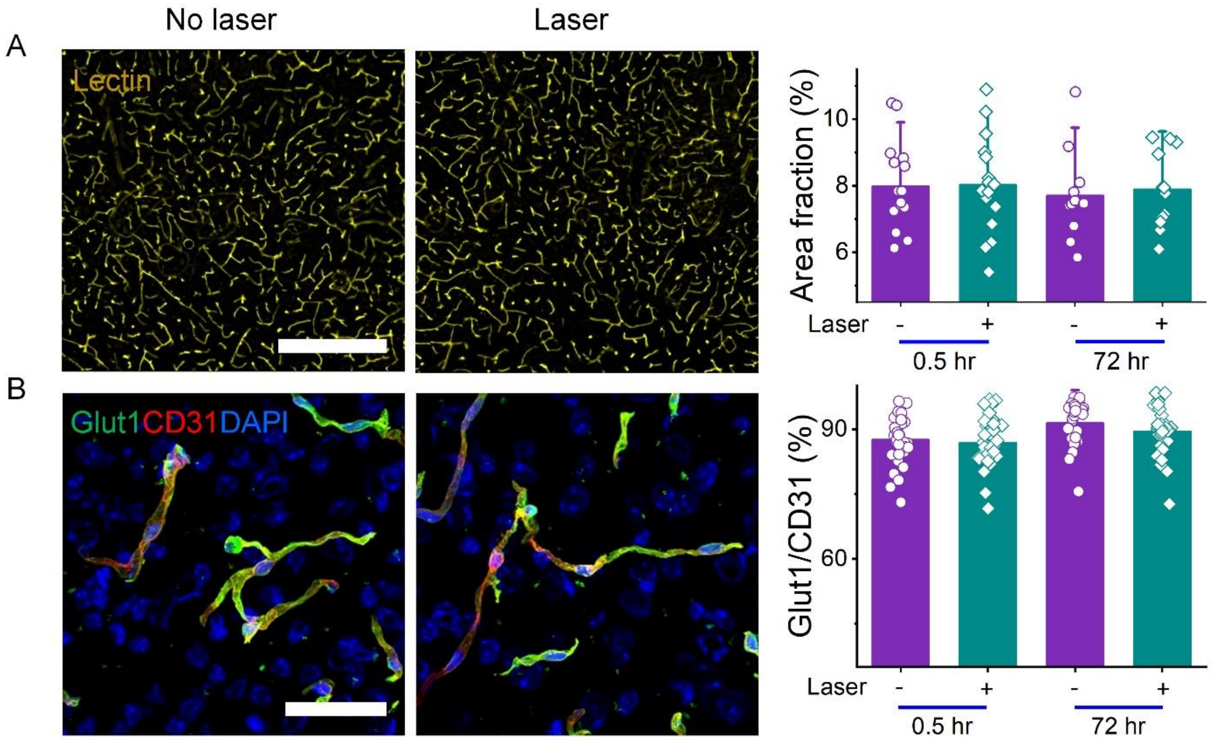


**Fig. S9. The BBB modulation does not cause changes in capillary density (lectin-labeled vessels), glucose transporters (Glut1) (25 mJ/cm$^2$, 1 pulse)**. (A) The blood vessels were labeled by tomato lectin for density analysis indicated by area fraction of lectin per field. The quantification suggests no significant change in vessel density suggesting normal structure of brain vasculature. (B) IHC staining of glucose transporters Glut1. The signal of Glut1, normalized by CD31 indicating blood vessels, shows no significant difference suggesting normal function of brain vasculature. Scale bar: 400 µm (A), 40 µm (B). Data expressed as Mean ± SD (n >10).

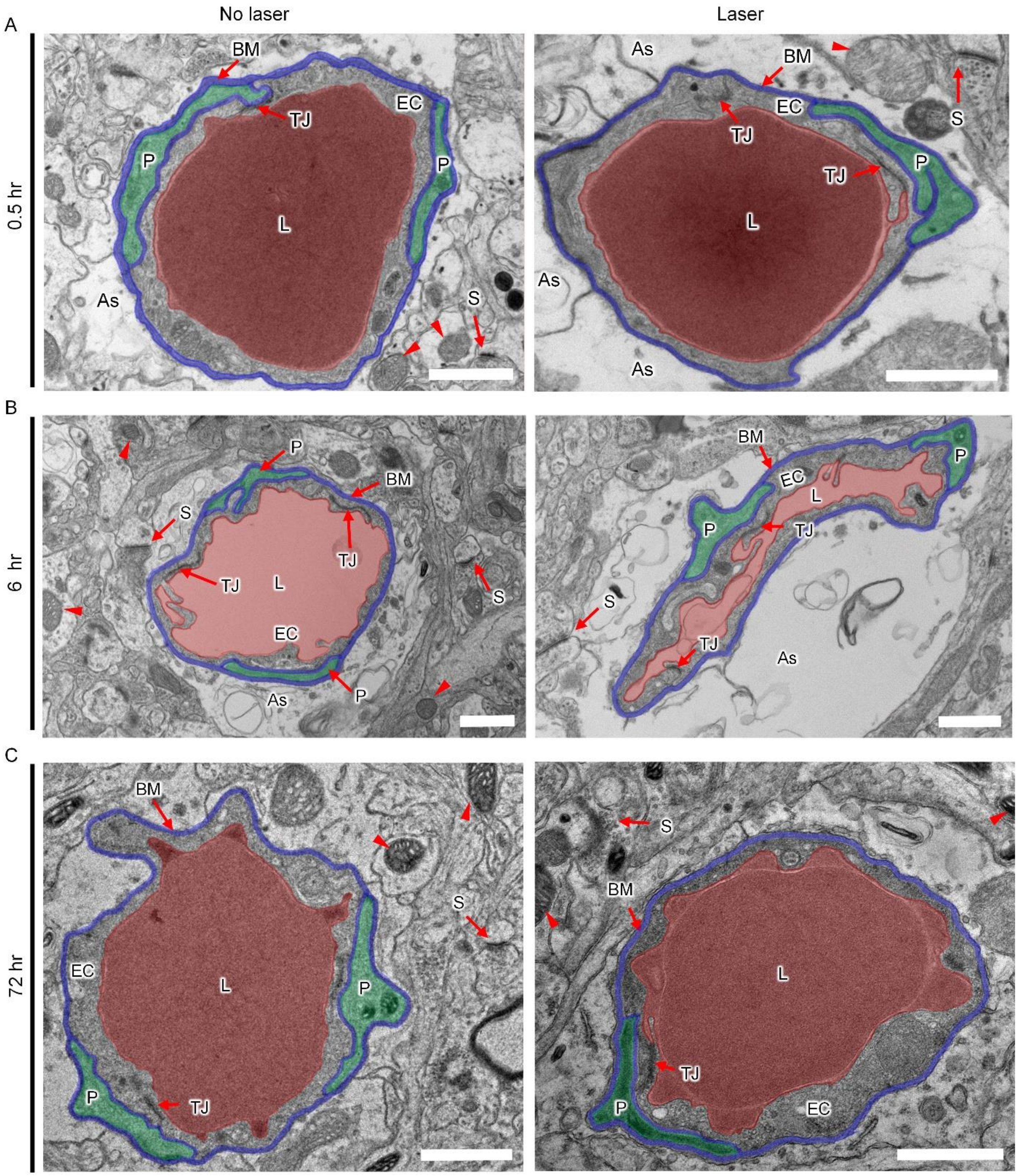


**Fig. S10. The brain ultrastructure under BBB modulation (5 mJ/cm$^2$, 1 pulse)**. The tight junction (TJ), pericyte (P), basement membrane (BM) in ipsilateral side appears intact at 0.5, 6 and 72 hours, compared with those in untreated contralateral side. The enlarged astrocyte endfeet are observed at 0.5 and 6 hours. The synapse and mitochondria appear no difference post laser exposure. Pseudocolours: Lumen (L: red), basement membrane (BM: blue), pericyte (P: green). Endothelial cell (EC), astrocyte endfeet (As), synapse (S), mitochondria (Arrow head). Scale bar: 1 μm.

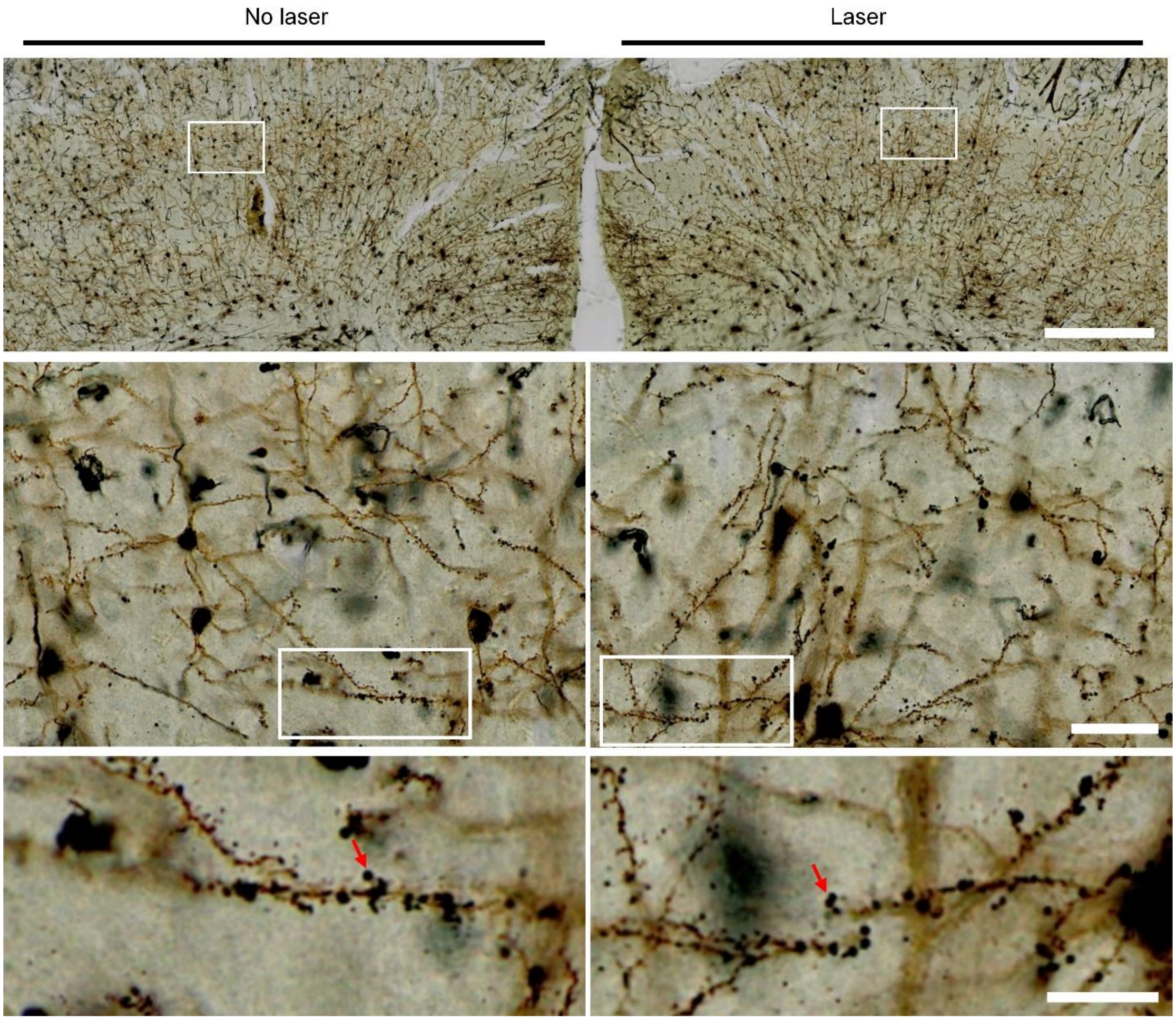


**Fig. S11. The Golgi staining shows optical BBB modulation preserves dendritic spines of neurons (25 mJ/cm$^2$, 1 pulse, 72 hours post laser stimulation).** Dendritic spines (arrow) appear the same with and without laser excitation. Scale bar, 500 µm (top panel), 50 µm (middle panels), 20 µm (bottom panels).

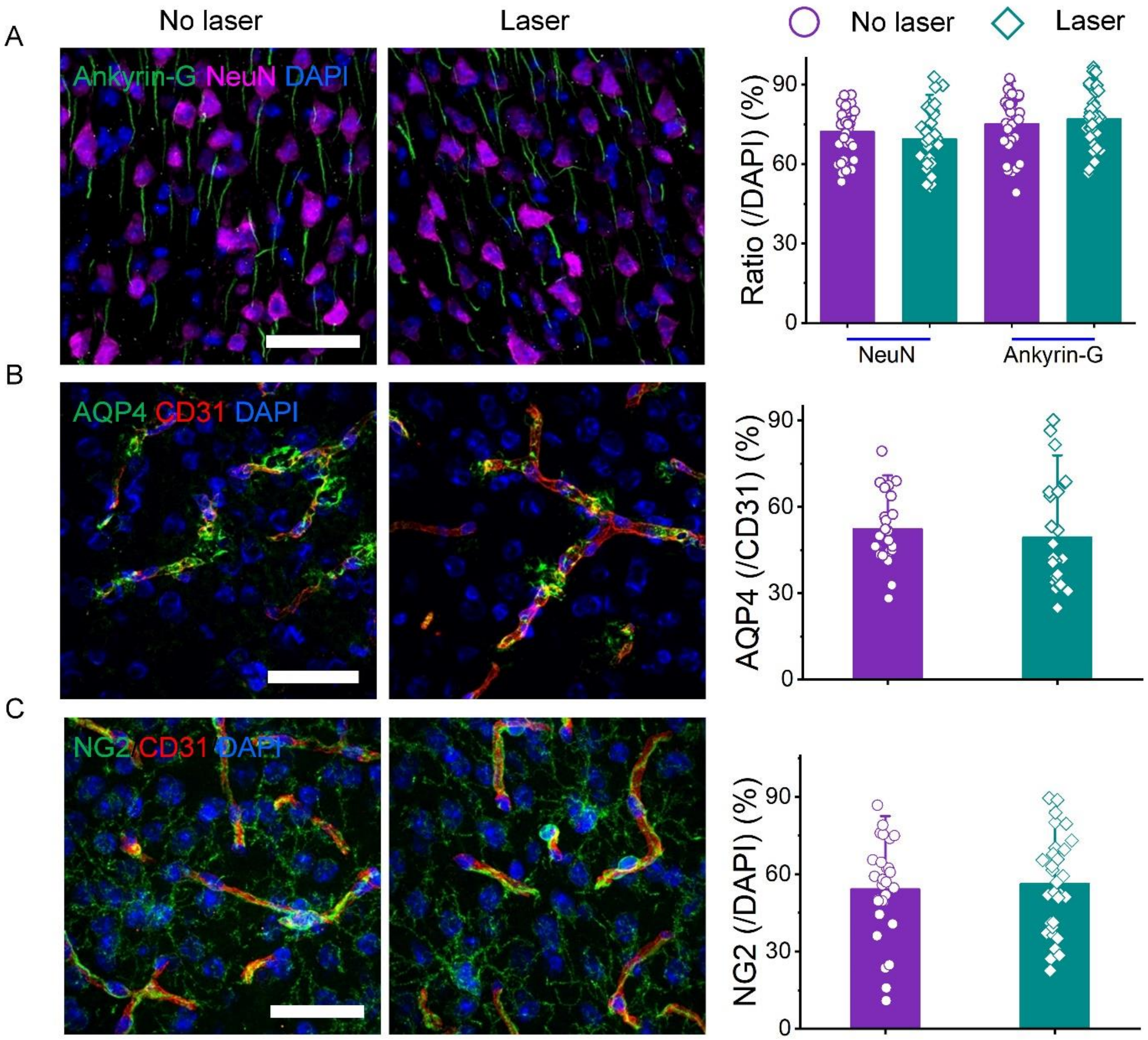


**Fig. S12. BBB modulation preserves the cellular architecture of the brain parenchyma (25 mJ/cm$^2$,1 pulse, 72 hours post laser stimulation).** The IHC staining of neuronal nucleus and axon indicated by NeuN and Ankyrin-G (A), water transporter of astrocyte endfeet indicated by AQP4 (B) and pericyte indicated by NG2. CD31 indicates blood vessels. The quantification analysis of NeuN, Ankyrin-G, AQP4 and NG2 shows there is no significant difference with and without laser treatment. Scale bar, 40 μm [(A), (B), (C)]. Data expressed as Mean ± SD (n≥20).

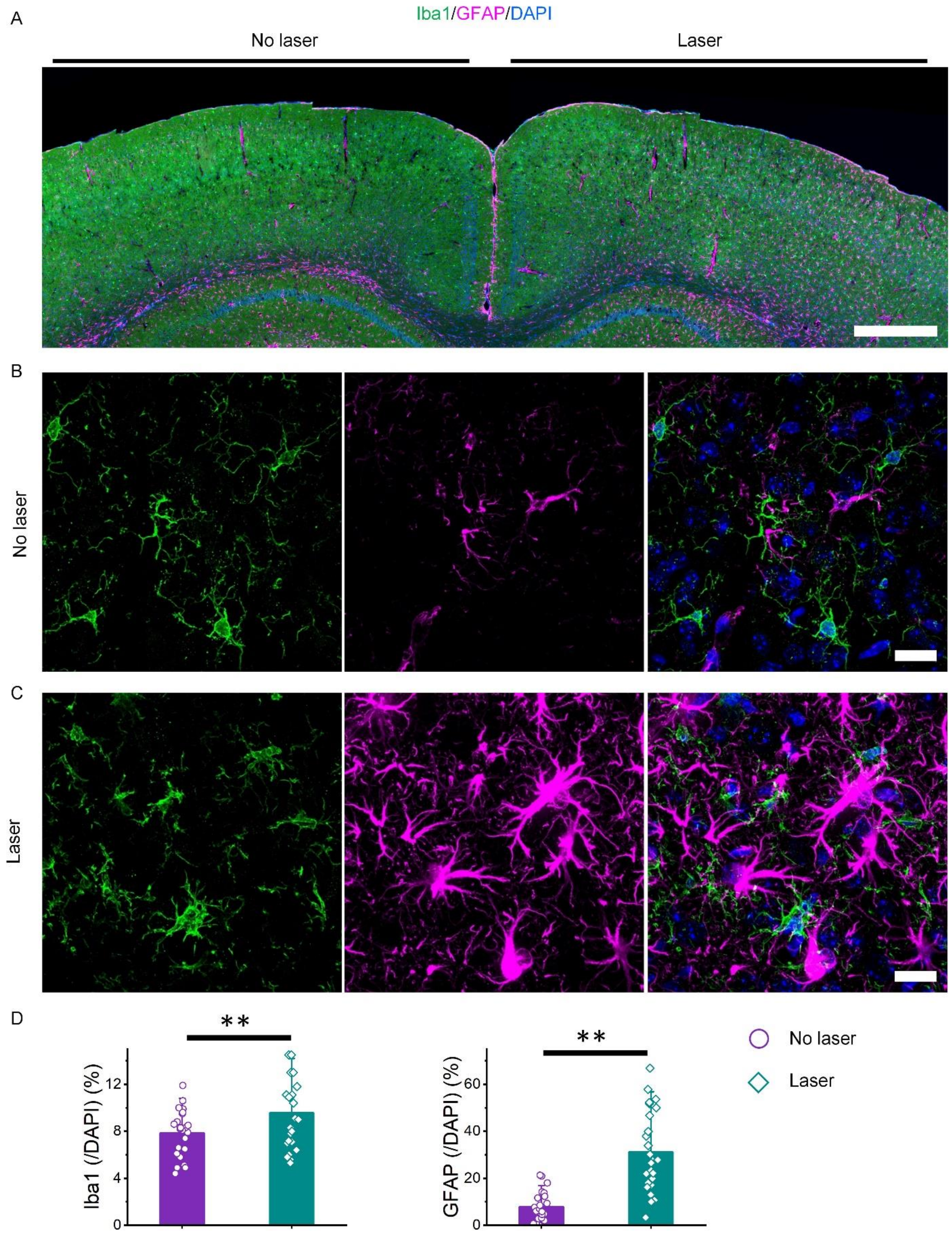


**Fig. S13. Microglia and astrocyte increase after laser treatment (25 mJ/cm$^2$,1 pulse, 72 hours post laser stimulation).** (A) Slide scanning image shows more microglial indicated by Iba1 and astrocytes indicated by GFAP at laser treatment region. (B) The confocal images show the Iba1$^+$

microglial and $GFAP^+$ astrocyte without laser (B) and with laser (C) treatment. (D) Quantification analysis of Iba1 and GFAP shows the $Iba1^+$ microglial and $GFAP^+$ astrocyte increased significantly. Scale bar, 500 μm (A), 20 μm [(B), (C)]. Data expressed as Mean ± SD (n ≥25). **P<0.05

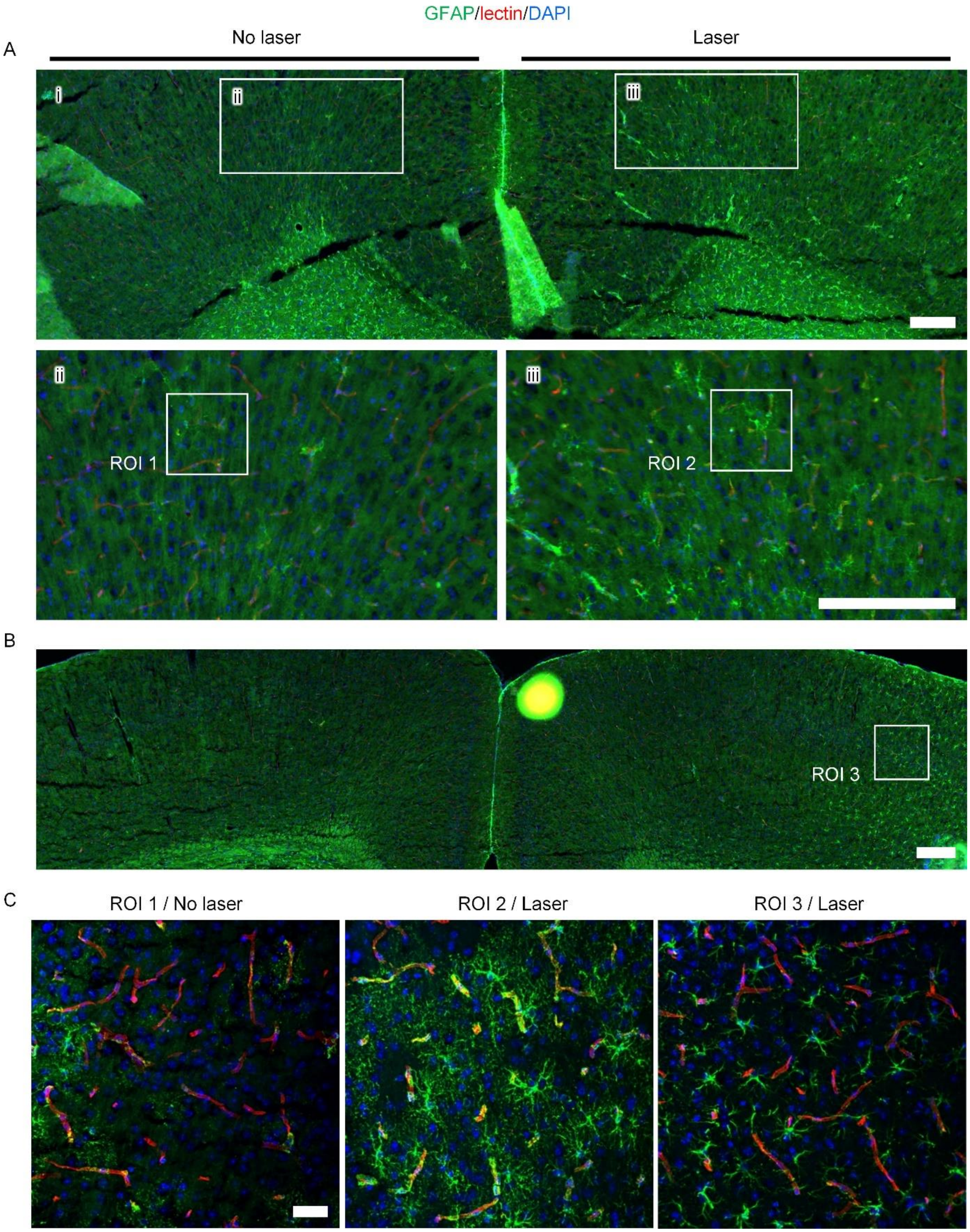


**Fig. S14. The increase of reactive astrocytes show heterogeneity after laser treatment (5 mJ/cm$^2$, 1 pulse, 72 hours post laser excitation)**. The IHC staining of reactive astrocyte indicated by GFAP. Slide scanning images show more reactive GFAP$^+$ astrocytes in cortex with laser treatment region [(A), (B)]. The confocal images of $GFAP^+$ astrocytes without laser treatment (C, ROI 1) and with laser treatment (C, ROI 2 and ROI 3). The reactive $GFAP^+$ astrocytes reveal various morphology change at different regions, which suggests the heterogeneity of reactive $GFAP^+$ astrocyte. Scale bar, 200 μm [(A), (B)], 40 μm (C).

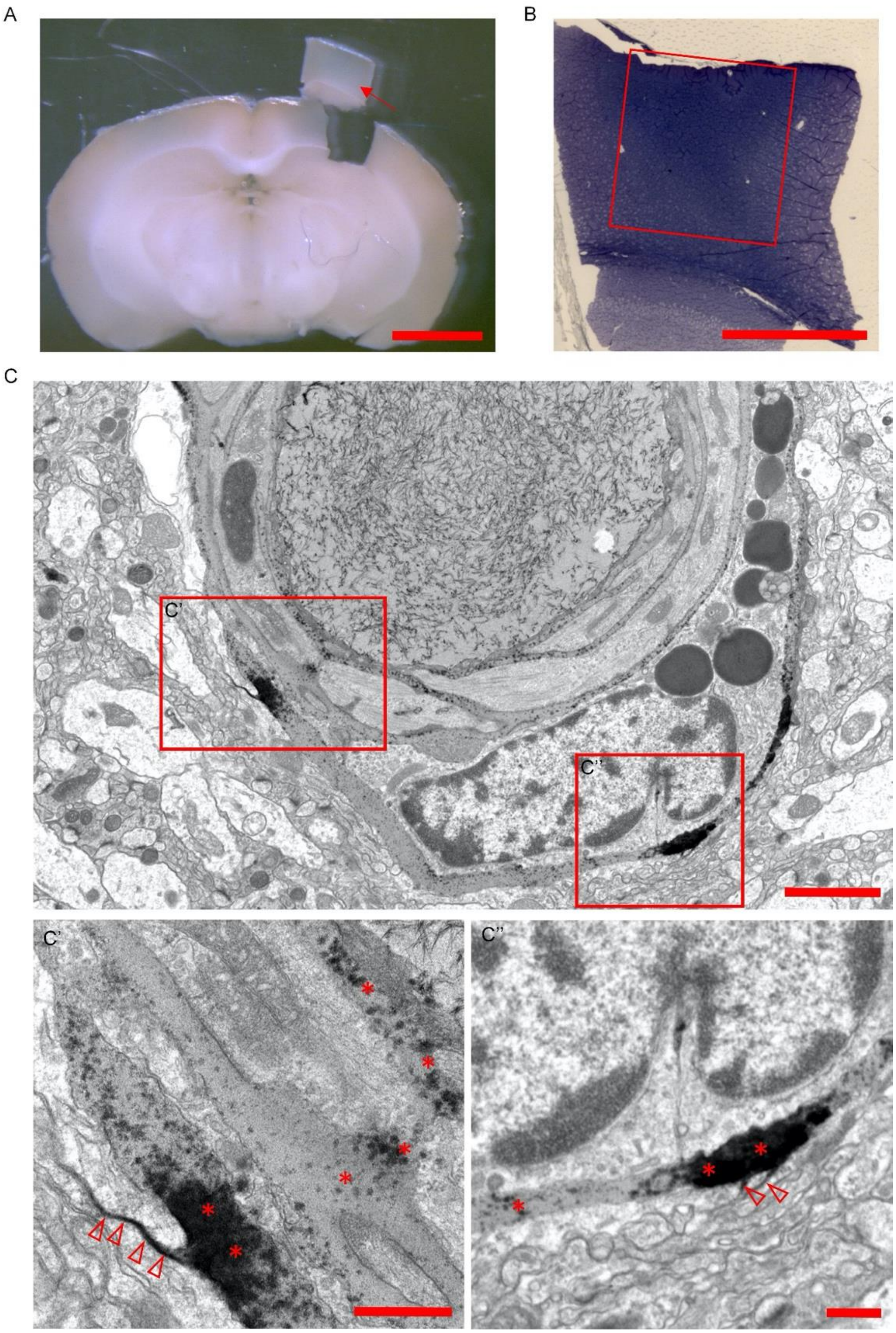


**Fig. S15. Lanthanum diffuse into basement membrane and interstitial space post laser excitation of TJ-targeted AuNPs (25 mJ/cm$^2$, 1 pulse, 2 hours post laser treatment).** (A) Using Evan blue as indicator to identify the BBB modulation area for further processing. Arrow: The selected region for preparation of EM sample. (B) The semi-thin section before processing for EM imaging. Red frame: the area for EM imaging. (C) Lanthanum diffusion into basement membrane

(*) and interstitial space (empty arrow head). Scale bar: 2 mm (A), 0.5 mm (B), 2 µm (C), 500 nm [(C'), (C'')].

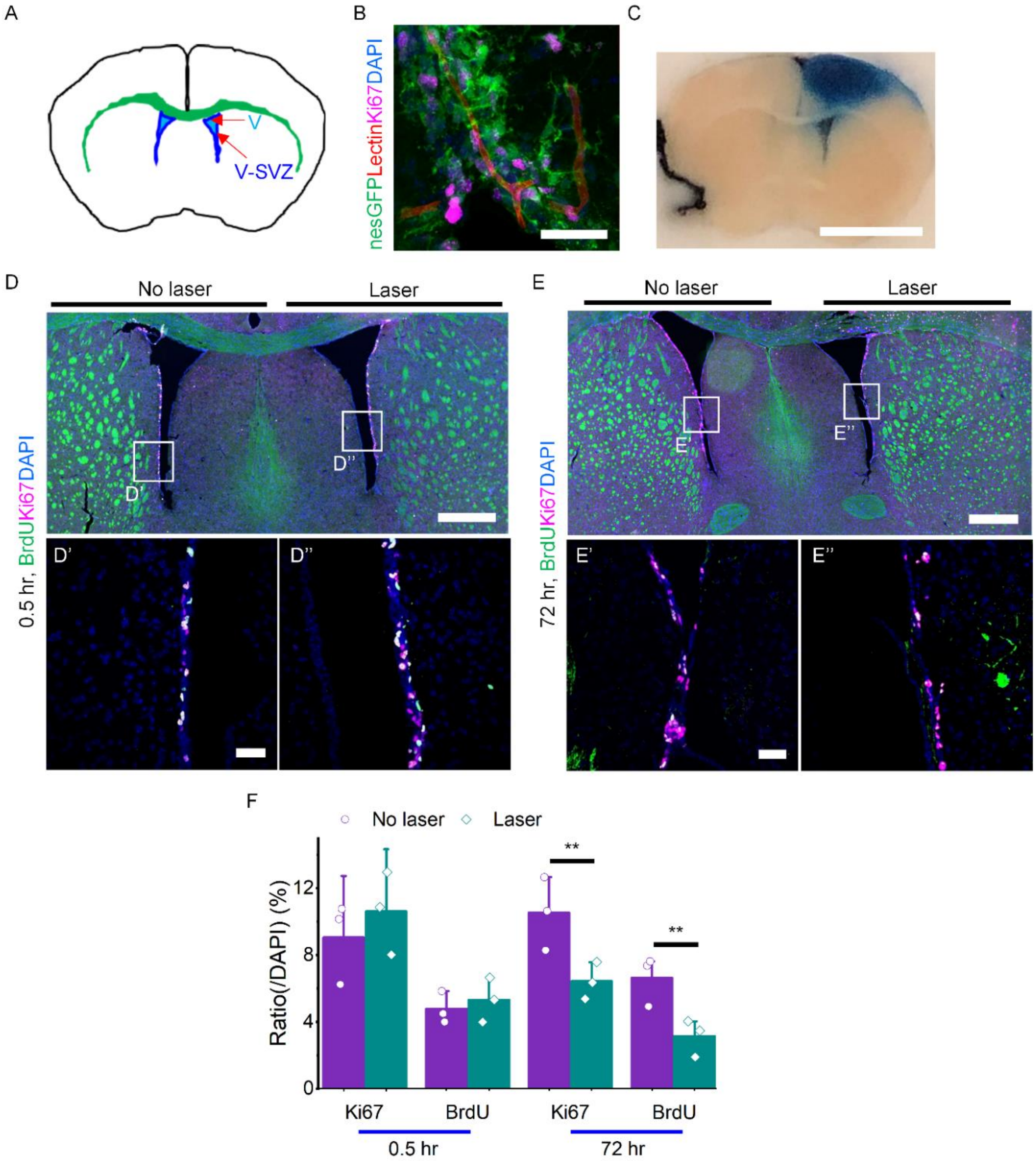


**Fig. S16. BBB modulation disrupts neural stem cell niche in ventricular-subventricular zone (V-SVZ).** (A) Schematic showing the location of V-SVZ. (B) Confocal image shows the proliferating neural stem cell in V-SVZ interacted intimately with blood vessel in nestin: GFP mice. Green: nestin-GFP indicates neural stem cells. Red: lectin labels the blood vessel. Magenta: ki67 marks the dividing cell. (C) BBB modulation in V-SVZ indicated by Evans blue signal (48 mJ/cm$^2$, 1 pulse). (D-E) IHC staining of neural stem cells in V-SVZ at 0.5 hours and 72 hours post laser excitation. Green: BrdU, Magenta: ki67, Blue: DAPI. (F) Quantification of dividing neural stem cell. Scale bar: 40 μm [(B), (D'), (D''), (E), (E'')], 4 mm (C), 500 μm [(D), (E)]. Data expressed as Mean ± SD (n=3). ** $p < 0.05$.

Table S1. The optical properties of mouse brain

| λ = 532 nm | | d [cm] | $\mu_a$ [cm$^{-1}$] | $\mu_s$ [cm$^{-1}$] | g | $\mu_s' = \mu_s(1-g)[cm^{-1}]$ | n | |
|---|---|---|---|---|---|---|---|---|
| | | Thickness | Absorption coefficient | Scattering coefficient | Anisotropy factor | Reduced scattering coefficient | Refractive index | Ref. |
| Skull bone | | 3.00E-02 | 13.6 | 351.4 | 0.93 | 24.6 | 1.5 | (*41*) |
| Meninges | CSF | 1.00E-02 | 0.004 | 2.5 | 0.001 | 2.5 | 1.33 | (*42*) (*43*) |
| Cerebral cortex | grey matter | 1.00E-01 | 0.44 | 100 | 0.88 | 12 | 1.3951 | (*44*) |
| | white matter | 3.00E-02 | 0.95 | 423 | 0.806 | 82.1 | 1.4121 | (*44*) |
